# The genetic architecture of floral trait divergence between hummingbird- and self-pollinated monkeyflower (*Mimulus*) species

**DOI:** 10.1101/2024.06.04.597389

**Authors:** Hongfei Chen, Colette S. Berg, Matthew Samuli, V. Alex Sotola, Andrea L. Sweigart, Yao-Wu Yuan, Lila Fishman

## Abstract

1. Pollination syndromes are a key component of flowering plant diversification, prompting questions about the architecture of single traits and genetic coordination among traits. Here, we investigate the genetics of extreme floral divergence between naturally hybridizing monkeyflowers *Mimulus parishii* (self-pollinated) and *M. cardinalis* (hummingbird-pollinated).
2. We mapped quantitative trait loci (QTLs) for 18 pigment, pollinator reward/handling, and dimensional traits in parallel sets of F_2_ hybrids plus recombinant inbred lines and generated nearly isogenic lines (NILs) for two dimensional traits, pistil length and corolla size.
3. Our multi-population approach revealed a highly polygenic basis (n = 190 QTLs total) for pollination syndrome divergence, capturing minor QTLs even for pigment traits with leading major loci. There was significant QTL overlap within pigment and dimensional categories. Nectar volume QTLs clustered with those for floral dimensions, suggesting a partially shared module. The NILs refined two pistil length QTLs, only one of which has tightly correlated effects on other dimensional traits.
4. An overall polygenic architecture of floral divergence is partially coordinated by genetic modules formed by linkage (pigments) and likely pleiotropy (dimensions plus nectar). This work illuminates pollinator syndrome evolution in a model radiation and generates a robust framework for molecular and ecological genomics.

## Introduction

Across flowering plants, distantly related taxa often show similarities in a suite of floral phenotypes that can be recognized as pollination syndromes (Fenster *et al*., 2004; Dellinger, 2020), while switches between pollination syndromes are common even among closely related species. For example, the evolution from bee-to hummingbird-pollination, which is characterized by red color, copious nectar, and stigma and anthers exerted beyond a large bill-accommodating corolla, has happened more than 10 times independently in *Penstemon* alone (Wilson *et al*., 2007; Wessinger & Hileman, 2016). Similarly, autogamous self-pollination, which is associated with inconspicuous coloration, reduced nectar rewards, and reduced anther-stigma separation (Sicard & Lenhard, 2011), has evolved countless times within animal-pollinated lineages (Stebbins, 1970; Barrett, 2002; Goodwillie *et al*., 2005). Both convergence and divergence in pollination syndromes requires the correlated evolution of multiple traits to maintain floral phenotypic integration and reproductive fitness throughout the entire evolutionary path. Three non-exclusive genetic mechanisms may contribute to such coordinated evolution of pollination syndromes and other complex multi-trait strategies. At one extreme, natural selection on floral traits may be strong enough to restrict successful plants to a few discrete adaptive peaks even in the face of gene flow (Bleiweiss, 2001), building stereotypical multi-trait pollination syndromes from variation at multiple unlinked loci (Wessinger *et al*., 2023). At the other extreme, pleiotropy among floral traits (Troth *et al*., 2018) may enforce coordinated evolution of trait modules during pollination syndrome divergence (Smith, 2016; Wessinger & Hileman, 2016). Finally, genome architectures that suppress recombination in heterozygotes can package genes for functionally distinct traits into adaptive supergenes (Lowry & Willis, 2010; Hermann *et al*., 2013; Edwards *et al*., 2021; Liang *et al*., 2023). Distinguishing among these explanations reveals very different barriers to traversing the phenotypic landscape as flowers evolve coordinately from one multi-phenotypic optimum to another.

Over the past three decades, quantitative trait locus (QTL) mapping has revealed the genetic architecture of pollination syndrome divergence between numerous closely-related pairs of plant species, including three sections of *Mimulus* monkeyflowers (Bradshaw *et al*., 1998; Fishman *et al*., 2002, 2013, 2015; Stankowski *et al*., 2023)*, Petunia* (Stuurman *et al*., 2004)*, Ipomoea* (Rifkin *et al*., 2021; Liao *et al*., 2021) and many others. Across angiosperm diversity from monocots to diverse eudicots, these genome-wide approaches reveal two broad patterns. First, divergence in pollinator-attraction and reward traits such as flower color, scent, or nectar volume is often controlled by few loci, each of moderate to large effect, whereas divergence in dimensional traits (corolla or reproductive organ size and shape) often involves more loci, each of small effect. This pattern may in part reflect focus on attraction/reward traits in studies of transitions among distinct animal pollination syndromes vs. a primary focus on floral dimensions in shifts to self-pollination. However, it may also reflect consistent differences in the underlying genetic variation and patterns of selection for different categories of trait. Second, co-localization of QTLs for at least some floral traits suggests floral integration/modularity through pleiotropy and/or the adaptive evolution of supergene architecture to facilitate trait packaging in the face of gene flow (Yeaman & Whitlock, 2011). Floral integration/modularity has been considered a plausible mechanism that facilitates rapid evolution of pollination syndromes (Diggle, 2014; Wessinger *et al*., 2014; Smith, 2016; Dellinger *et al*., 2019; Kostyun *et al*., 2019). However, few studies have clearly identified intra-floral evolutionary modules, and the pattern of integration and modularity across sets of floral traits remains an open question. Understanding both the build-up (integration) and the breakdown (modularity) of trait correlations to generate complex floral strategies requires a locus-and gene-scale understanding of the genomic bases of pollination syndromes.

Although QTL mapping has robustly advanced understanding of the genetic architecture of divergence in multi-factorial trait syndromes, connecting their effects to the underlying genes remains a challenge. First, especially when traits are polygenic and the underlying loci have small effects, scans of single experimental hybrid mapping populations capture the overall architecture of genetic variation, but not individual loci and their effects. Thus, replication of mapping experiments tests QTL robustness to environmental variation and increases confidence in shared QTLs. Second, even major QTLs often contain 10s to 100s of genes, confounding pleiotropy and linkage as causes of genetic correlation and QTL coincidence. Similarly, fine-scale identification of causal variants benefits from isolation from segregating background effects. Construction of near-isogenic lines (NILs) allows both detection and isolation, as in the identification of major flower color and scent loci in *Petunia* (e.g. (Klahre *et al*., 2011; Berardi *et al*., 2021) and *Mimulus* (Bradshaw & Schemske, 2003; Yuan *et al*., 2013b; Byers *et al*., 2014; Yuan *et al*., 2016; Liang *et al*., 2023). However, because NIL construction generally involves strong selection for resemblance to the introgressing parent for a focal trait combined with opposing selection on other traits, it may not capture the evolutionary contributions of more complex genetic architectures to trait (co-)variation. Thus, a combination of genome-wide QTL characterization plus targeted NIL construction is a powerful approach to understand both the genome-wide architecture and gene-scale causes of divergence in complex pollination syndromes.

Here, we employ an integrated approach to characterize the genetic architecture of pollination syndrome divergence and the detailed genetics of key traits between closely related monkeyflowers *Mimulus cardinalis* and *M. parishii* (Phyrmaceae, section *Erythranthe*). These taxa, which are hummingbird-pollinated and self-pollinated, respectively (Fig. 1), are the most florally divergent members of a recent adaptive radiation (Nelson *et al*., 2021b). Nevertheless, they hybridize in areas of range overlap, producing genomic signatures of recent introgression (Nelson *et al*., 2021b). Along with bee-pollinated *M. lewisii*, which likely resembles the common ancestor of all three taxa, these closely related species are a model system for understanding the genetics of floral divergence and speciation (Yuan, 2019). Work in this system has provided key insights into the evolution and molecular biology of divergent pollination syndromes and their roles in local adaptation and speciation (Hiesey *et al*., 1971; Bradshaw *et al*., 1995; Schemske & Bradshaw, 1999; Ramsey *et al*., 2003; Fishman *et al*., 2013; Yuan *et al*., 2013b, 2014; Stathos & Fishman, 2014; Fishman *et al*., 2015; Yuan *et al*., 2016; Peng *et al*., 2017; Nelson *et al*., 2021b,a; Liang *et al*., 2023). New resources, including chromosome-scale reference genomes (www.Mimubase.org), dense linkage maps (Sotola *et al*., 2023), and stable transformation protocols (Yuan, 2019), now enable genome-wide mapping, gene-scale dissection, and molecular characterization of the loci underlying *Erythranthe* floral diversity. Notably, because *Mimulus parishii* x *M. cardinalis* hybrids segregate more freely than other crosses in the group (Fishman *et al*., 2013, 2015; Sotola *et al*., 2023), they were key to the recent genetic and molecular dissection of a novel speciation supergene (Liang *et al*., 2023). With the current study, we take a major step toward a similarly detailed understanding of the full suite of floral traits contributing to pollination syndrome divergence.

**Fig. 1.**
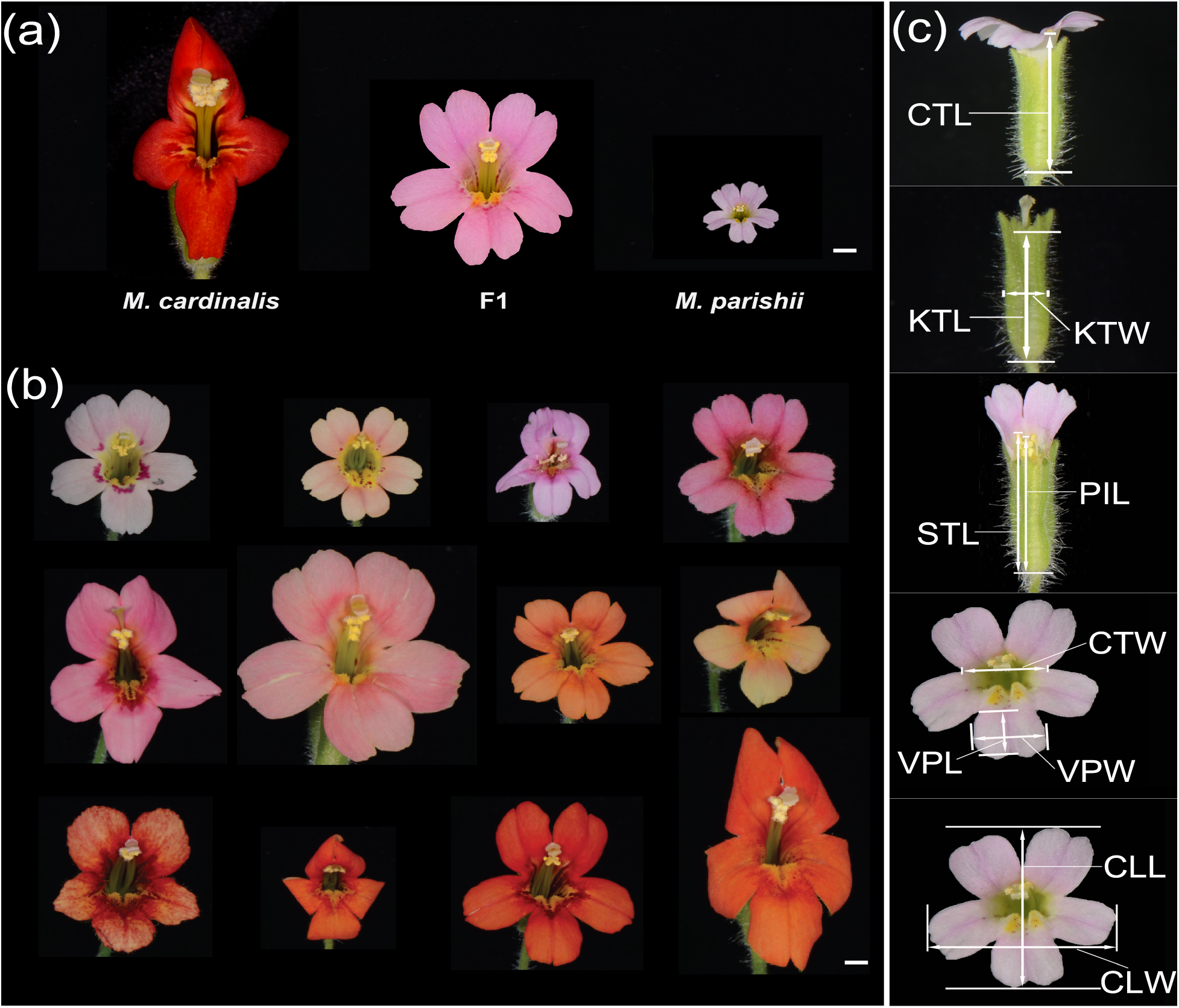
Floral phenotypes. (a) The parental line *M. cardinalis* CE10, *M. parishii* PAR, and their F1 hybrid. (b) Representative F2 progeny. (c) The main floral traits measured in this study, using *M. parishii* as examples. CTL: corolla tube length; KTL: calyx tube length; KTW: calyx tube width; STL: stamen length; PIL: pistil length; CTW: corolla tube width; VPL: ventral petal length; VPW: ventral petal width; CLL: corolla limb length; CLW: corolla limb width. Scale bars, 3 mm.

To robustly characterize the genetic architecture of extreme floral divergence, we examine patterns of floral trait (co)inheritance and map QTLs in one extensively phenotyped focal F_2_ population, then map QTLs for a subset of traits in independent F_2_ and RIL growouts to capture additional minor QTLs in distinct environmental and genetic backgrounds. We assess patterns of QTL size and coincidence within and across trait categories and propose three hypotheses about the genetic architecture: (i): Divergence of floral traits produced by relatively simple biochemical pathways such as flower color are controlled by few loci with moderate to large effects, whereas dimensional traits involve more loci of small effects; (ii) minor QTLs are more subject to stochasticity and differences in environment and genetic background (for RILs) among our mapping populations; and (iii) traits within a category (e.g., pigment or dimensions) are controlled by integrated sets of overlapping QTLs (modules) while overlap between categories is relatively low. Finally, we characterize independent nearly isogenic lines (NILs) for two highly polygenic dimensional traits, flower size and pistil length; these NILs provide proof of concept (and sound some useful cautionary notes) on the dissection of floral dimensional QTLs. Our multi-trait and multi-generation approach provides a broad and deep characterization of the genetics of floral divergence and opens paths toward understanding both its molecular bases and effects on patterns of mating and introgression in wild populations.

## Materials and Methods

### Study system, mapping populations, and phenotyping

Hummingbird-pollinated *M. cardinalis* (Phrymaceae) is a perennial herb of low-elevation seeps and riverbanks from northern Baja California to southern Oregon (Angert & Schemske, 2005; Angert, 2009). It has red flowers with long tubular corolla, copious nectar, and exserted stigma and anthers (Fig. 1a). *M. parishii* is an annual self-pollinating herb generally found along ephemeral streams in southern California. *M. parishii* has small pale pink flowers, little stigma–anther separation, and no nectar (Fig. 1a, Table 1). All hybrids in this study were generated from two highly inbred parental lines: Sierran CE10 for *M. cardinalis* (Yuan *et al*., 2013b) and PAR for *M. parishii* (Fishman *et al*., 2015; Nelson *et al*., 2021b; Liang *et al*., 2023). F_1_ hybrids were generated with PAR as the seed parent and selfed to generate F_2_ seeds, while recombinant inbred lines (RILs) were generated by single-seed-descent from F_2_ individuals through 3-6 generations of self-fertilization (Sotola *et al*., 2023). F_2_ hybrids (Fig. 1b) were grown in two separate greenhouse common gardens at the University of Connecticut (UC_F_2_) and the University of Montana (UM_F_2_), and the RILs were grown at the University of Georgia (UGA), as detailed in Supplementary Methods S1.

**Table 1.** Floral trait variation (means +/- SE for *M. cardinalis* (CE10), *M. parishii* (PAR), and their F_1_ and F_2_ hybrids in UC_F_2_ growout. Broad-sense heritability (*H^2^*) was calculated following Fishman *et al*. (2002). Traits marked ** were also measured in both UM_F_2_ and RIL mapping populations, while nectar volume (NEV; *) was also characterized in RILs.

| Trait | *M. cardinalis*<br>CE10 (n=8) | *M. parishii*<br>PAR (n=8) | F<sub>1</sub> hybrid<br>(n=8) | F<sub>2</sub> hybrids<br>(n=253) | $H^2$ |
| --- | --- | --- | --- | --- | --- |
| **Pigment** |  |  |  |  |  |
| Petal lobe carotenoid (PLC)** | 99.38 ± 0.44 | 2.29 ± 0.50 | 13.89 ± 0.48 | 48.01 ± 2.12 | 1.00 |
| Petal lobe anthocyanin (PLA)** | 74.39 ± 1.11 | 21.46 ± 1.93 | 64.15 ± 1.02 | 52.38 ± 1.23 | 0.96 |
| Nectar guide anthocyanin (NGA) | 9 | 1 | 5 | 5.20 ± 0.12 | NA |
| **Pollinator reward/handling** |  |  |  |  |  |
| Nectar volume (NEV)* | 45.95 ± 3.83 | 0 ± 0 | 3.28 ± 0.17 | 5.30 ± 0.29 | -0.35 |
| Nectar guide trichome (NGT) | 7 | 1 | 4 | 4.40 ± 0.10 | NA |
| **Dimension** |  |  |  |  |  |
| Corolla limb length (CLL) | 36.45 ± 0.69 | 8.18 ± 0.09 | 19.97 ± 0.49 | 20.21 ± 0.26 | 0.89 |
| Corolla limb width (CLW) ** | 20.23 ± 1.62 | 8.94 ± 0.09 | 18.61 ± 0.43 | 18.30 ± 0.22 | 0.49 |
| Ventral petal width (VPW) | 12.46 ± 0.39 | 3.28 ± 0.07 | 7.27 ± 0.12 | 7.52 ± 0.09 | 0.82 |
| Ventral petal length (VPL) | 10.43 ± 0.21 | 2.77 ± 0.06 | 5.51 ± 0.10 | 6.11 ± 0.07 | 0.91 |
| Corolla tube width (CTW)** | 8.29 ± 0.27 | 4.43 ± 0.27 | 7.69 ± 0.15 | 7.26 ± 0.08 | 0.84 |
| Calyx tube length (KTL) | 22.30 ± 0.17 | 9.82 ± 0.28 | 16.17 ± 0.29 | 16.90 ± 0.14 | 0.90 |
| Calyx tube width (KTW) | 7.68 ± 0.13 | 2.98 ± 0.08 | 5.06 ± 0.08 | 5.34 ± 0.06 | 0.91 |
| Corolla tube length (CTL)** | 32.85 ± 0.35 | 11.74 ± 0.31 | 21.68 ± 0.23 | 22.46 ± 0.20 | 0.93 |
| Pistil length (PIL)** | 45.88 ± 0.43 | 11.76 ± 0.21 | 25.89 ± 0.29 | 27.20 ± 0.28 | 0.96 |
| Stamen length (STL)** | 42.93 ± 0.28 | 13.31 ± 0.47 | 24.91 ± 0.27 | 25.30 ± 0.24 | 0.94 |
| Stigma-anther separation (SAS)** | 2.95 ± 0.28 | -1.55 ± 0.34 | 0.99 ± 0.16 | 1.90 ± 0.09 | 0.71 |
| **Flowering time (FLT)\*\*** | 79.38 ± 0.84 | 53.5 ± 1.07 | 55.13 ± 0.30 | 66.68 ± 0.51 | 0.94 |

In the UC_F_2_ growout, we measured three pigment, two pollinator reward/handling and nine dimensional floral traits, plus flowering time, on F_2_s (n = 253) plus parental lines and F_1_ hybrids (n =8 each) (Fig. 1). We scanned the ventral petal of each flower to quantify petal lobe anthocyanin (PLA) and carotenoid (PLC) pigment intensity. The proportion of red (R), green (G), and blue (B) pixels in a square area of the same size of the adaxial surface of the petal were estimated from scanned images using Image J (http://rsbweb.nih.gov/ij/). The relative petal lobe anthocyanin concentration was estimated using the equation “[(R + B)/2] − G”, a simple and effective approach previously used for genetic mapping of anthocyanin content variation (Yuan *et al*., 2013b) and independently verified in other plant species (Valle *et al*., 2018). Similarly, the relative carotenoid concentration was estimated by the equation “[(R + G)/2] – B”, which also proved effective as our QTL mapping successfully located the previously characterized carotenoid locus *YELLOW UPPER* (*YUP*) (see Results). Nectar guide anthocyanin (NGA) and nectar guide trichome length (NGT) were visually scored in F_1_ and F_2_ hybrids on semi-quantitative scales defined by the parental extremes (CARD = 9 and 7, respectively, PAR = 1 for both). Nectar volume (NEV) was measured for two flowers per individual on their first day of opening, using a pipette accurate to 1.5μL. To reduce environmental effects, nectar was measured at 4:00 PM-7:00 PM after watering at 12:00-1:00PM each day. Floral dimensions (Fig. 1c) were measured on one of the second pair of open flowers using a digital caliper, and stigma-anther distance calculated as pistil length – stamen length. F_2_ trait distributions were tested for normality with a Shapiro–Wilk test implemented in R v. 3.6.0. Because some traits were non-normally distributed, we calculated pairwise Spearman’s correlation coefficients (r) for phenotypic correlations using the *psych::corr.test* function in R v. 3.6.0; we calculated broad-sense heritability for each trait and genotypic correlations following (Fishman *et al*., 2002).

Overlapping subsets of key floral traits were measured using parallel methods in the UM_F_2_ (n = 278) and UGA_RIL growouts (n = 145) (Supplementary Methods S2). Traits with a shared abbreviation represent the same floral dimension, except for pistil length (PIL) and stigma-anther separation (SAS), which included the stigma lobes in the UC_F_2_ and UGA_RILs (Fig. 1c) but not the UM_F_2_s. In the UM_F_2_ growout, we characterized an additional pollinator handling trait associated with pollination syndrome divergence in *Mimulus*, touch-sensitive stigma closure (Friedman *et al*., 2017). The bilobed stigmas of *M. cardinalis* close rapidly (<5s; like a tiny venus flytrap) when touched, while *M. parishii* stigmas are insensitive and/or non-closing (Fishman *et al*., 2024). Prior to the other floral measurements, a single tester touched each stigma head-on with a pencil eraser to mimic pollinator contact and scored stigma closure speed on a 4-point scale (0 = no closure = PAR-like, 3 = fast closure = CE10-like, 1 and 2 = slower and faster intermediates, respectively).

### Genetic context and QTL mapping

We previously constructed a joint linkage map of the two F_2_ populations and a separate map of the RILs using windowed genotypes from ddRAD sequences aligned to the CE10 *M. cardinalis* reference genome (Sotola *et al*., 2023). The dense F_2_ and RIL linkage maps are generally highly collinear with each other and the physical genome assemblies (www.Mimubase.org). However, an *M. cardinalis*-specific reciprocal translocation involving portions of Chromosomes 6 and 7 (Fishman *et al*., 2013; Stathos & Fishman, 2014) causes inter-chromosome linkage (i.e. they form a single linkage group in F_2_s: LG6&7), excess heterozygosity, and underdominant hybrid sterility. In addition, a gametophytic Dobzhansky-Muller incompatibility involving Chr4 (∼7-8 Mb) and Chr8 (∼12-40 Mb) eliminates three genotypic classes in F_2_ and later hybrids (Sotola *et al*., 2023).

We conducted QTL mapping in QTL Cartographer (Wang *et al*., 2005) in parallel on the three populations using composite interval mapping (model 6, with forward-backward regression to choose 10 cofactors, window size 10 cM). LOD significance thresholds for QTL detection for each trait were set with 1000 permutations. Due to substantial retained heterozygosity in the RILs (Sotola *et al*., 2023), we used F_2_ rather than RIL population settings to allow full estimation of QTL effects using all individuals. To evaluate QTL coincidence across populations and traits, as well as define physical bounds, we defined a 1.5 LOD-drop confidence interval (CI) around each peak. For the two F_2_ populations, which share a linkage map, overlap was directly determined. For comparing F_2_s and RILs, we translated QTL peaks and intervals to the physical positions of boundary markers. We assigned QTL numbers within traits across populations based on CI overlap (Table S1). We tested whether the mean effect size (r^2^) of all 144 unique (at level of trait) QTLs differed among the trait categories using ANOVA in JMP 18. For the nine traits measured in all three populations (94 named QTLs), we similarly tested whether QTLs detected in one (n = 55), two (n = 32) or three (n = 7) populations were, on average, of different magnitude. Using the physical positions of all QTLs in Table S1, we calculated the degree of QTL overlap between each pair using the Jaccard index, following (Liao *et al*., 2021) (Supplementary Methods S3), then calculated the mean (and standard error) of QTL overlap within and between trait categories. We tested whether there was greater overlap within than between trait categories using 1000 permutations in which traits were randomly assigned to categories. Because some non-dimensional traits may plausibly share a partial genetic basis with flower size, we also specifically assessed the overlap of nectar volume and flowering time QTLs with those in the three multi-trait categories.

### Construction and characterization of nearly isogenic lines (NILs)

NILs were constructed via phenotypic selection prior to QTL mapping, and thus provide an independent approach to dissecting the genetics of dimensional traits. To construct pistil length (PIL) NILs in the *M. parishii* genetic background, we chose an F_2_ individual with overall similarity to *M. parishii* in both floral and vegetative traits, but with conspicuously longer pistil, for serial backcrossing (with NIL as pollen donor) to *M. parishii.* From each backcross growout of ∼95 plants, we selected one individual that closely resembled *M. parishii* but with longer pistil for the next round. Bulked segregant analysis in a BC_2_S_1_ population (two rounds of backcrosses followed by one round of selfing) and subsequent genotyping in the same population using markers within the identified fragments revealed two chromosomal regions (Chr 4: 0-4.1 Mb; Chr 6: 42-52 Mb) introgressed from *M. cardinalis* that co-segregate with pistil length. Further genotyping of a BC_3_S_1_ population narrowed the chromosome 6 locus to a genomic interval at 42.85 Mb-51.76 Mb (Supplementary Methods S4). Selfing a BC_3_S_1_ individual heterozygous for both fragments generated nine genotypes across the two loci, which also allowed us to decompose the BC_3_ NIL into two NILs. A similar crossing approach was used to generate a corolla limb length (CLL) NIL representing flower size (Supplementary Methods S4).

## Results

### Floral trait divergence, distributions, and correlations in F_2_ hybrids

In the focal UC_F_2_ grow-out, the CE10 *M. cardinalis* and PAR *M. parishii* parental lines were highly differentiated for all traits, with F_1_ and F_2_ means always intermediate (Table 1). Floral dimensions were normally distributed in F_2_s, but petal carotenoid values (PLC) were bimodally distributed while petal lobe anthocyanins (PLA) were skewed toward CE10 and nectar volume (NEV) toward PAR (Fig. S1).

Quantitative traits other than NEV, which had negative H^2^ estimates due to its extreme skew, had high broad-sense heritability (H^2^ > 0.49). The UM_F_2_ and RILs had similar distributions for each shared trait, but mean nectar volume was much higher in the RILs (Fig. S1). All floral dimensions other than stigma-anther separation (SAS) were positively correlated both phenotypically (r_P_) and genetically (r_G_) (Fig. S2). The key mating system trait of stigma-anther separation was most highly correlated with pistil length (r_G_ = 0.57), less with the other length metrics (r_G_ = 0.25 - 0.31), and uncorrelated with width metrics. Floral dimensional traits were only moderately correlated with flowering time (FLT) but flower length traits (KTL, CTL, STL, PIL) were highly correlated with nectar volume (r_P_= 0.58-0.70, r_G_ not calculable for NEV due to negative H^2^) and petal lobe carotenoids were strongly correlated with pistil length and stamen length (both r_G_ > 0.6).

### Genetic architecture - QTL mapping of individual traits in multiple mapping populations

We identified 190 floral QTLs, which define 144 QTL locations if collapsed (within traits) across the three mapping populations (Fig. 2, Table S1).

**Fig. 2.**
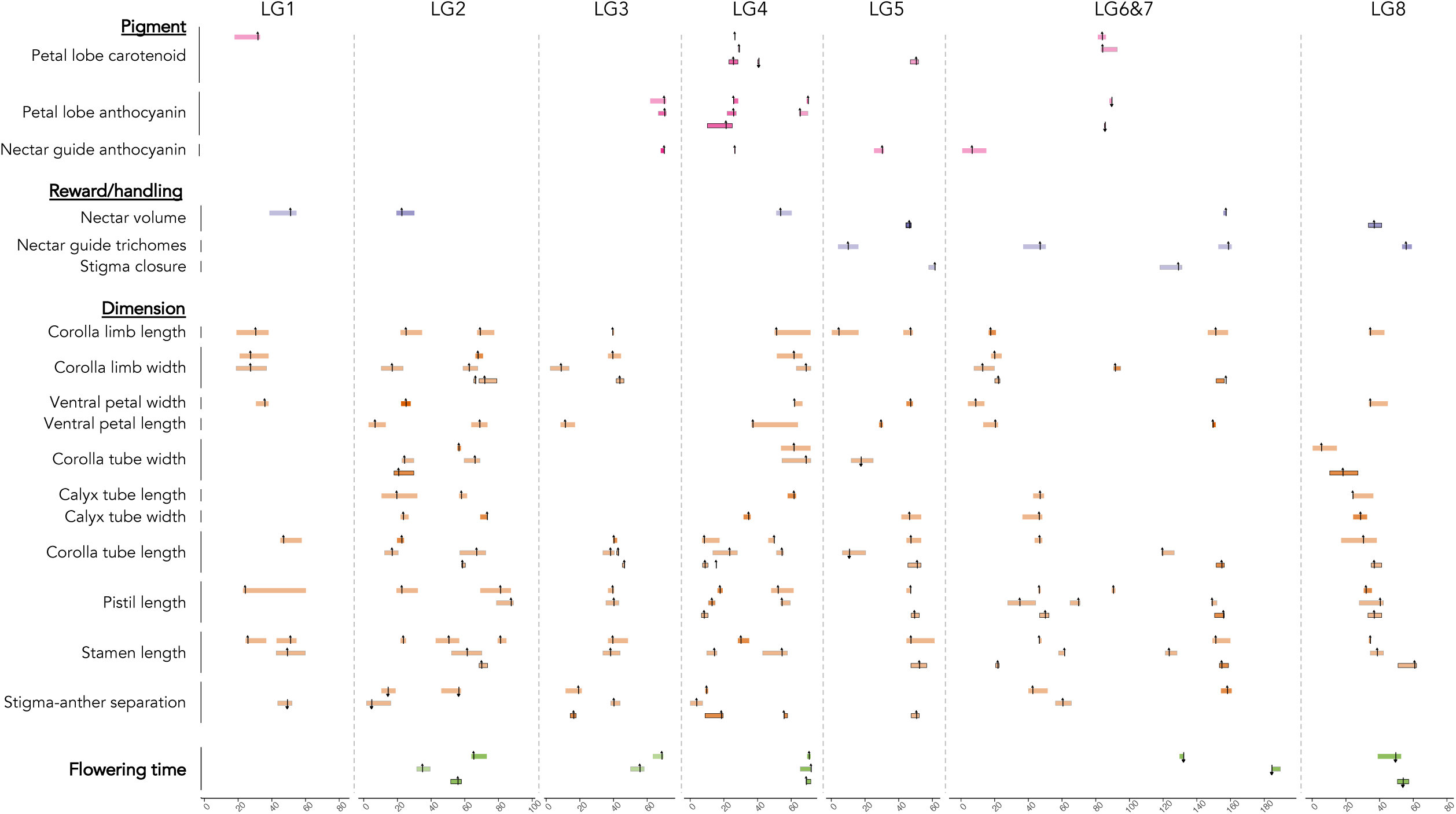
Quantitative trait loci (QTLs) for floral traits associated with pollination syndrome divergence between selfer *Mimulus parishii* and hummingbird pollinated *M. cardinalis*, as detected in three mapping populations. Bars show QTL 1.5 LOD drop confidence intervals and arrows indicate QTL peak position (up = QTL effect is in direction expected from parental divergence, down = opposite). The UC_F_2_ QTL bars (all traits other than stigma closure) are unbordered, UM_F_2_ QTL bars are bordered in gray, and RIL QTL bars are bordered in black. Relative QTL magnitude is indicated by the color-intensity of the QTL bar. The x-axis is position in centiMorgans (cM) on each of the seven F_2_ linkage groups; these correspond to eight chromosomes due to a reciprocal translocation between Chr 6 and Chr 7 in *M. cardinalis* vs. *M. parishii* that generates inter-chromosomal linkage (Fishman *et al*., 2013; Stathos & Fishman, 2014; Sotola *et al*., 2023)

#### Pigment traits

As expected from previous work, petal lobe carotenoids (PLC) were primarily affected by a fully shared *M. cardinalis*-recessive major QTL on LG4 (coincident with *YUP*; PLC4.1 in Table S1). We also detected two smaller carotenoid loci in the F_2_s, and two more in the RILs. For petal lobe anthocyanins (PLA), four loci were detected: PLA4.1 was detected in all three growouts and coincident with the *YUP-SOLAR-PELAN* supergene, PLA4.2 and PLA3 were found in both F_2_s but not the RILs, and PLA6&7 was found in both RILs and UC_F_2._ Nectar guide anthocyanins (NGA) were under the control of two major loci, NGA3 (r^2^ = 0.21) and NGA4 (r^2^ = 0.31), and two additional small QTLs.

#### Reward and handling traits

We detected four nectar volume QTLs in the UC_F_2_ and two completely non-overlapping ones in the RILs. RIL QTLs NEV5 and NEV8 had absolutely ∼4x larger effects than the largest F_2_ one (NEV6&7; r^2^ = 0.20), which explained only ∼1/7 of the parental difference in NEV. The four largest NEV QTLs (RIL pair, plus NEV2 and NEV6&7) were each coincident with dense clusters of floral dimension QTLs (see below). Nectar guide trichomes (NGT) and stigma closure speed (SCS) QTLs, which were scored on semi-quantitative scales, had relatively low explanatory power in the segregating F_2_ populations (all r^2^: 0.04-0.11). However, QTLs for these traits explained from 20% (each of the two *M. parishii*-recessive stigma closure QTLs: SCS5 and SCS6&7) to 40% of the parental difference (additive NGT8). Thus, they provide key targets for understanding the genetic underpinnings of these important but understudied components of floral syndrome evolution.

#### Dimensional traits and flowering time

In the UC_F_2_, floral size was polygenic (71 dimensional QTLs). QTL sizes were correspondingly small, with the leading QTL for each trait explaining from 11% (CLL, CTL, KTW) to ∼20% (VPL, VPW) of the F_2_ variance. All primary size QTLs in this F_2_ population moved trait values in the direction expected from the parental difference. Consistent with the transgressive segregation of stigma-anther separation in the UC_F_2_s (Fig. S1), 2 of the 6 QTLs for this composite trait had opposite effects from expectation (Table S1). For the six shared dimensional traits, we mapped 43, 39, and 29 QTLs in the UC_F_2_, UM_F_2_ and RILs, respectively, and ∼1/3 (35/111) were shared across two or more populations. For flowering time, FLT4.1 (all three mapping populations) and FLT8.1 (UC_F_2_ and RILs) were shared, but the other 7 QTLs were each found in only a single population. Flowering time QTLs are not particularly small in absolute terms (all 2*a* > 6 days), so this variation may reflect true genotype x environment interactions for phenology.

### Patterns across trait categories – genetic architecture, repeatability, modularity, and directionality

Overall, QTLs for pigment traits were nearly twice as large as dimensional QTLs (0.135 vs.0.075, p = 0.003), while flowering time and handling/reward QTLs were intermediate (Fig. 3a). For shared traits, pigment QTLs and larger ones were significantly more likely to be detected in all three mapping populations (both P < 0.005). However, QTLs detected in one or two populations were equally small (0.074 vs. 0.077, P = 0.94 by Tukey’s HSD). This pattern of moderate repeatability suggests that each mapping population stochastically detected only a subset of minor loci from the larger (shared) pool of variants influencing each polygenic trait.

**Fig. 3.**
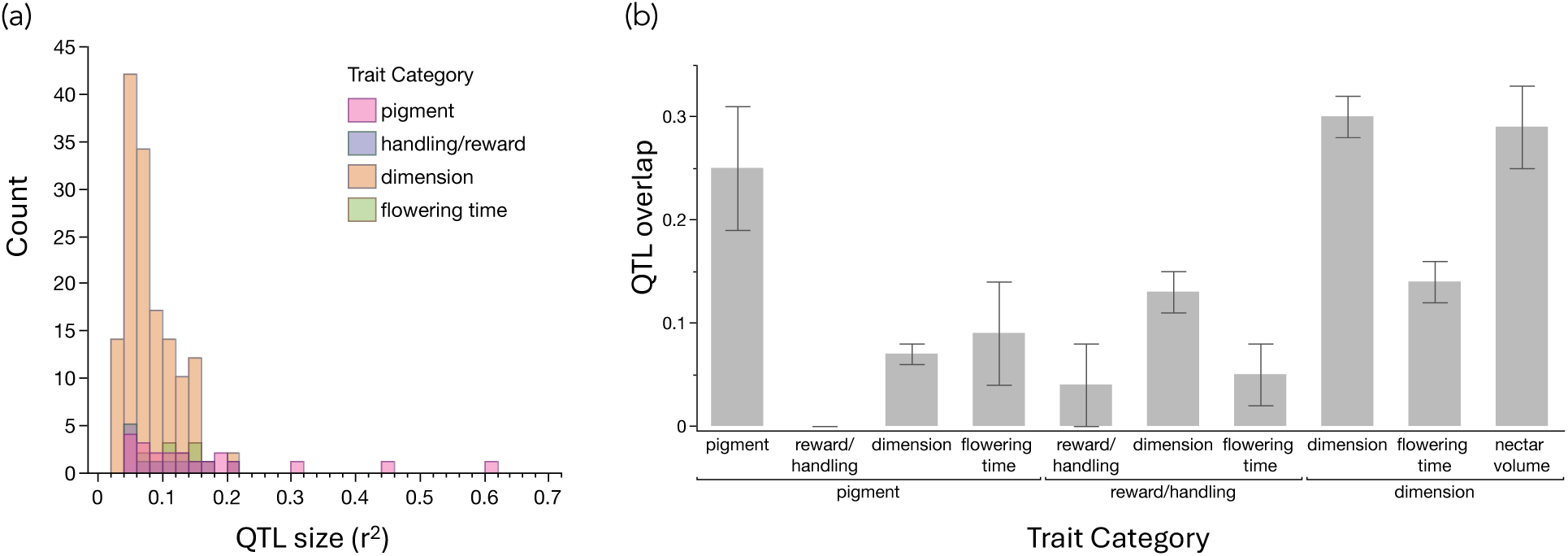
Summary of QTL effects by trait category. (a) Effect size (r2) for QTLs in pigment, pollinator reward/handling, and dimensional categories, plus flowering time. (b) QTL overlap (Jaccard index ± 1SE) within and between the three multi-trait categories, plus overlap of each with nectar volume and flowering time.

Overlap of QTLs within both pigment (Jaccard index = 0.25) and dimension (0.30) categories were significantly greater than null expectation (p = 0.036 and 0.0001, respectively), suggesting that each forms a distinct intra-floral evolutionary module (Fig. 3b). Much lower overlap (0.04) within the pollinator reward/ handling set is not surprising, given its grab-bag of traits. However, nectar volume QTLs strongly overlapped with those for floral dimensions (0.29), suggesting a joint evolutionary module with flower size, while overlap of flowering time and dimension QTLs was intermediate (Fig. 3b). Overall, only 7% (10/144) unique QTLs had additive effects opposite to those expected from the parental difference, suggesting consistent divergent natural selection (Orr, 1998). Notable exceptions were flowering time and the composite floral trait of stigma-anther separation, with >1/4 and 1/3 (respectively) of their QTLs opposite to expectation.

### Dissection of floral dimension QTLs with NILs

The long-pistil (PIL) and flower size (CLL) NILs isolated *M. cardinalis* alleles in the *M. parishii* background using repeated rounds of backcrossing and selfing with phenotypic selection (Supplemental Methods S3). The long-pistil NIL contained two unlinked regions introgressed from *M*. *cardinalis*, corresponding to QTLs *PIL4.1* (0-4.1 Mb on Chr4) and *PIL6&7.2* (42.85 Mb-51.76 Mb on Chr6) (Fig. 4). In a BC_3_S_1_ population segregating for all genotypic combinations at these two loci, both QTLs exhibited primarily additive effects on pistil length, consistent with QTL effects. The two NIL- isolated QTLs both had absolute effects smaller than in the F2 population but maintained their relative sizes: *PIL4.1* (2*a* = 5.68mm in F2s, 3mm in NIL) vs. *PIL6&7.2* (2*a* = 4.24mm in F2s, 1mm in NIL). There was no evidence of an epistatic interaction between these two loci, as the double NIL was 4mm larger than *M. parishii* (Table S2). The CE10-homozygous *PIL4.1* NIL also increased stamen and corolla tube lengths, but not our measure of overall flower size (CLL), relative to *M*. *parishii* (Table 2). In contrast, the *PIL6&7.2* NIL had little effect on stamen or corolla tube length (Table S2), suggesting that it contains a gene that specifically regulates pistil length.

**Fig. 4.**
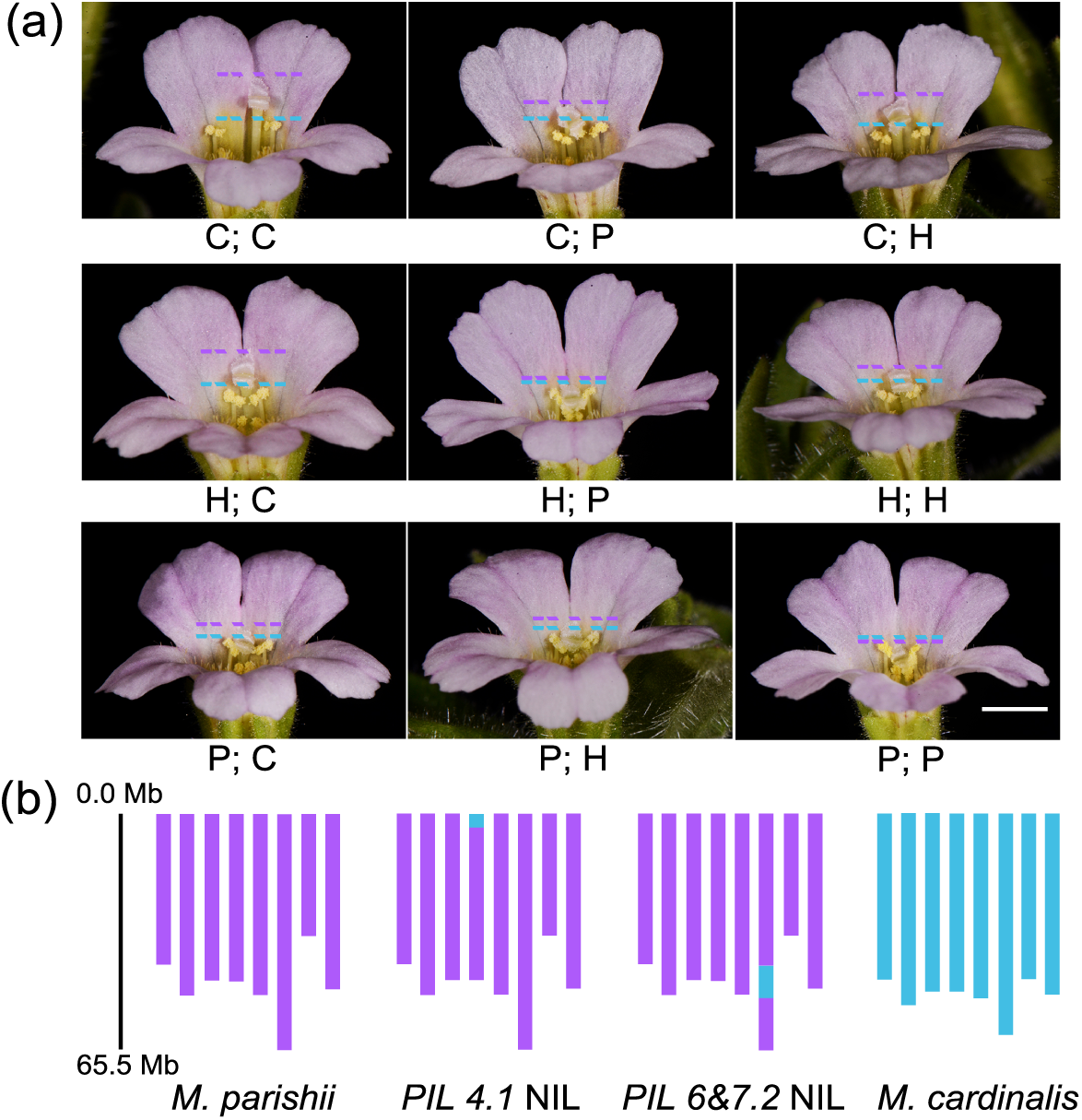
Pistil NILs. (a) Phenotypes of 9 genotypic combinations of the *PIL4.1* and *PIL6&7.2* loci. Genotypic designations: C, homozygous for *M. cardinalis*; H, heterozygous; P, homozygous for *M. parishii*. Under each image, the *PIL4.1* genotype is followed by the *PIL6&7.2* genotype. The colored dashed lines indicate the positions of the stigmas and long anthers, respectively. Scale bar, 3 mm. (b) Graphical genotypes of the *PIL4.1* NIL and the *PIL6&7.2* NIL. The colored bars represent the 8 chromosomes.

After selfing and phenotypic selection (Supplemental Methods S3), the final BC_3_S_1_ NIL for corolla limb length (CLL) captured a large region including *CLL6&7.2* (Fig. 5). This NIL was heterozygous across much of Chr6 (43.54Mb-60.3Mb) and Chr7 (0-29.34Mb), but *M. cardinalis* homozygous near the LG6&7 translocation breakpoint (60.32Mb-61.12Mb on Chr6; 29.34Mb-29.82Mb on Chr7). Unlike the more tightly localized *PIL6&7.2* introgression (which it includes in heterozygous state), the *CLL6&7.2* NIL has increased pistil, stamen, and corolla tube length, as well as greater CLL, relative to *M. parishii* (Fig. 5a, Table S3). Because the *CLL6&7.2* NIL includes a recombination-suppressed translocation breakpoint associated with underdominant pollen sterility (Sotola *et al*., 2023), its multiple phenotypic effects may reflect *PIL6&7.2*, additional linked genes, and pleiotropic effects of sterility per se (Fishman et al. 2015).

**Fig. 5.**
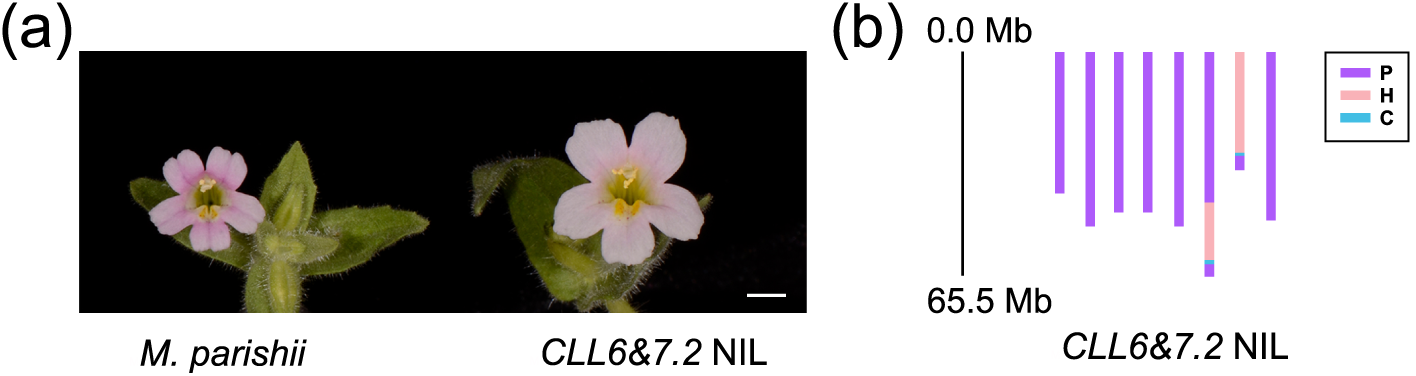
Flower size NIL. (a) Flowers of *M. parishii* and the *CLL6&7.2* NIL. Scale bar, 4mm. (b) Graphical genotype of the *CLL6&7.2* NIL.

## Discussion

We used QTL mapping in three hybrid growouts, as well as NIL construction, to investigate the genetic architecture of pollination syndrome divergence between hummingbird-pollinated *Mimulus cardinalis* and self-pollinated *M. parishii.* Despite some post-mating barriers (Sotola *et al*. 2023), these taxa are good models for understanding the early stages of speciation: hybrids between *M. parishii* and *M. cardinalis* are florally diverse (Fig. 1), more fit than their respective hybrids with bee-pollinated *M. lewisii* (Fishman *et al*., 2013, 2015; Stathos & Fishman, 2014), and subject to ongoing introgression in areas of range overlap (Nelson *et al*., 2021b). Along with directly illuminating the quantitative genetic basis of floral evolution, this work provides a foundation for understanding the molecular genetic basis, evolutionary history, and speciation consequences of complex pollination syndromes.

Overall, we identified a notably complex genetic basis for divergence between multi-trait pollination syndromes. Our findings largely confirm the initial hypotheses of a highly polygenic and thus less repeatably-mapped genetic basis for dimensional traits relative to pigment traits. Further, elevated genetic integration (shared QTL positions) within both pigment and dimensional trait categories (compared to low QTL overlap across categories) suggests trait modularity, as predicted. However, we also mapped new minor QTLs even for pigment traits influenced by known major supergenes (Liang *et al*., 2023), and characterized a complex (and background/environment-dependent) genetic basis for the key reward trait of nectar volume. This genetic architecture contrasts with both the highly oligogenic and tightly integrated (including across pigment, reward, and dimension traits) genetic architecture of divergence between *M. cardinalis* and bee-pollinated *M. lewisii* (Bradshaw *et al*., 1995, 1998; Fishman *et al*., 2013) and highly polygenic dimensional divergence between *M. parishii* and *M. lewisii* (Fishman *et al*., 2015)or selfer *M. nasutus* and bee-pollinated *M. guttatus* (Fishman *et al*., 2002). Along with chromosome-scale genomes and new functional genetic tools (Yuan, 2019), this complexity and diversity of genetic architectures reinforces the value of the *M. cardinalis* species complex for understanding the ecological, molecular, and evolutionary mechanisms of floral syndrome divergence.

### A long walk to dramatic floral divergence – genetic basis of individual traits

Although they show evidence of hybridization and ongoing introgression in areas of current range overlap (Nelson *et al*., 2021b), hummingbird pollinated *M. cardinalis* and selfer *M. parishii* likely both evolved from a bee-pollinated *M. lewisii*-like ancestor, which florally resembles their F_1_ hybrid (Fig. 1). Thus, it is not surprising that the genetic architecture of their floral divergence is a composite of patterns and loci seen in hybrids of each crossed to *M. lewisii.* Furthermore, the overall patterns of QTL size and directionality for different trait categories largely parallel predictions based on previous empirical work, as well as the underlying molecular and developmental pathways, where known. Specifically, we expected major leading QTLs for pigment traits and the reward trait of nectar volume. These key pollination syndrome traits may offer a limited set of mutational targets and pathways to adaptive evolution for both biochemical and evolutionary reasons (Bleiweiss, 2001; Wessinger & Rausher, 2012) and color divergence often maps to major on-off switches (Wessinger & Hileman, 2020). In contrast, dimensional traits may present near-infinite molecular targets for minor-effect mutations and exhibit high levels of intra-specific standing variation readily available for rapid polygenic adaptation under novel directional selection (Roels & Kelly, 2011; Troth *et al*., 2018). However, as summarized below, we find minor QTLs for all categories of traits in this wide cross. Although strong bias of QTL effects suggests that directional natural selection has driven trait divergence, this abundance of genetic contributors to variation in hybrids may reflect drift and relaxed selection along the lineage leading to selfer *M. parishii* compared to the more constrained shift between discrete bee- and hummingbird- attraction peaks (Bleiweiss 2001).

Consistent with expectation and previous genetics in this system, carotenoid (PLC) and anthocyanin (PLA, NGA) pigment traits each had a leading major QTL (r^2^ > 0.19) in the core UC_F_2_ population. Co-localized *PLC4.1* and *PLA4.1* correspond to the small RNA locus *YELLOW UPPER* (*YUP*) and the *R2R3-MYB* gene *PETAL LOBE ANTHOCYANIN* (*PELAN*), respectively (Liang *et al*., 2023), which form a pigment supergene novel to *Mimulus* section *Erythranthe* (Liang *et al*., 2023) along with another anthocyanin-regulating *MYB, SISTER OF LIGHT AREAS (SOLAR)* (Liang *et al*., 2022, 2023). It is unknown whether *PELAN* or *SOLAR* (or another linked gene) underlies the coincident nectar guide anthocyanin QTL (*NGA4.1*); nonetheless, QTL co-localization here reflects remarkably tight linkage of carotenoid and anthocyanin pigment variants affecting distinct pathways rather than pleiotropic effects of a single gene.

Beyond the *YUP-SOLAR-PELAN* supergene, however, our multiple maps revealed an unexpectedly complex and novel genetic basis for pigment traits. The four additional QTLs for each pigment trait had widely variable effects (Table S1), and only partially overlapped with *Mimulus* pigment loci identified with alternative approaches. For example, NGA3/PLA3 contains *RED TONGUE* (*RTO*), an R3-MYB gene previously shown to repress anthocyanin biosynthesis in both petal lobes and nectar guides of *M. lewisii* flowers (Ding *et al*., 2020). However, *ROSE INTENSITY (ROI)* (Yuan *et al*., 2013b), which controls the reduced anthocyanin of pale pink Sierran *M. lewisii* relative to *M. cardinalis*, was not coincident with any QTLs in this study, though PLA6&7, a rare wrong-way QTL at which *M. parishii* alleles confer darker anthocyanin pigmentation, contains several *R3-MYBs* related to *ROI* and *RTO*. Thus, there are clearly a diversity of mutational paths to complex floral pigment patterns in *Mimulus* flowers, paralleling the layers of complexity of similar pollination syndrome shifts in *Petunia* (Berardi *et al*., 2021). Overall, our pigment QTL data provide an unusually nuanced picture of the divergence of floral attraction traits, a roadmap for molecular characterization of the underlying genes, and the opportunity to study their effects on pollination ecology in natural and artificial hybrids.

The pollinator reward and handling traits (nectar volume, stigma closure speed, nectar guide trichomes) are each essentially lost in selfer *M. parishii* but may have followed distinct evolutionary paths to that endpoint. High nectar volume (NEV) maintains high hummingbird visitation rates to *M. cardinalis* (Schemske & Bradshaw, 1999), while touch-sensitive stigma closure (SCS) enhances pollen export in outcrossing monkeyflowers (Fetscher, 2001) and repeatedly degenerates in selfers (Friedman *et al*., 2017; Fishman *et al*., 2024). In contrast, nectar guide trichomes (NGT) may be under relaxed selection in both *M. cardinalis* and *M. parishii* relative to bee-pollinated *M. lewisii* (Chen & Yuan, 2024) and the common ancestor of these taxa. Like pigment traits, both nectar volume and nectar guide trichomes are under the control of major leading QTLs in *M. lewisii* x *M. cardinalis* and *M. parishii* x *M. lewisii* hybrids respectively, while loss of stigma closure appears moderately polygenic in yellow monkeyflowers (Fishman et al. 2024). Thus, QTLs for these traits, although not as extensively studied as floral pigments or dimensions, provide key comparative insight into the genetic architecture and (eventually) molecular basis of pollinator syndrome divergence.

Both nectar guide trichomes (NGT; UC_F_2_ only) and stigma closure (SCS; UM_F_2_ only) exhibit genetic novelty and complexity relative to parallel studies in *Mimulus*. Intriguingly, none of the four moderate NGT QTLs identified here includes the MYB transcription factor GUIDELESS (1.13 Mb on CE10 Chr6 (Chen & Yuan, 2024), previously inferred to be the major locus causal of NGT loss in *M. parishii* relative to bee-pollinated *M. lewisii* (Chen & Yuan, 2024) and also implicated as key determinant of nectar guide formation (both pigment and cell shape) in *M. lewisii* via mutagenesis (Yuan *et al*., 2013a). This suggests both a much more complex genetic basis for nectar guide divergence between these two non-bee species and possibly epistatic interactions masking nonsynonymous mutations inferred as causal of NGT loss in *M. parishii* (Chen & Yuan, 2024). Similarly, the two QTLs of moderate effect (SCS5 and SCS6&7) for stigma closure (which together explain only 1/3 of total F_2_ variance, suggesting many additional minor loci) do not map to chromosomal regions syntenic with the five QTLs that fully explain similar shift between selfer/noncloser *Mimulus nasutus* and fast-closer *M. guttatus* (Fishman *et al*., 2024). However, SCS6&7 contains a Mechanosensitive Channel of Small Conductance-like 10 (MSL10) gene homologous to a highly stigma-expressed candidate mechanosensor identified in the *M. guttatus* complex (Fishman *et al*., 2024). Furthermore, as in the *M. guttatus* complex, stigma closure QTLs appear independent of the other floral trait reductions associated with the evolution of selfing (Fig. 2, Table S1), suggesting abundant genomic targets for independent losses of this plant movement trait in selfers. Although pollinator-handling traits have not been as extensively studied as pigments, dimensions, or rewards, both nectar guide cell shape (e.g. (Glover *et al*., 1998) and stigma movement (Newcombe, 1922; Fishman *et al*., 2024) vary widely across Lamiales (>25,000 species). By revealing a diversity of underlying genetic mechanisms, even just within monkeyflowers, this work is a key step toward understanding the integration (and dis-integration in selfers) of pollination syndromes in diverse taxa with tubular, bilaterally symmetric flowers.

Nectar volume, with six small-to-moderate sized QTLs in the UC_F_2_s and RILs together, has an even more complex architecture, including polygenicity, epistasis, and gene x environment interactions. This sharply contrasts with the simple genetic architecture invoked for this trait in *M. lewisii* x *M. cardinalis* hybrids, which identified two leading QTLs each >30% of the F_2_ variance (Bradshaw *et al*., 1998). Our largest F_2_ QTLs, NEV2 and NEV6&7, were much smaller (r^2^ = 0.12-0.17) and all four summed to only ∼35% of the parental difference. Moreover, a model including all 2-way QTL interactions found significant (P < 0.005) interactions of NEV6&7 with NEV4 and NEV2; along with the strong skew of NEV phenotypes toward low (*M. parishii*-like) values (Fig. S1), this suggests that epistatic interactions may contribute to parental divergence. The genetics of nectar volume was also context-dependent; the two RIL QTLs were both much larger (10.9-14.3 µLs each) than any found in UC_F_2_s and in distinct locations. Differences between greenhouse conditions, as well as postzygotic barriers that further skew allele frequencies in RIL populations (Sotola *et al*., 2023), may contribute to the lack of repeatability. However, as discussed further below, nectar volume may be a particularly complex composite trait whose divergence encompasses both highly polygenic dimensional traits and simpler biochemical switches.

### Causes and consequences of floral integration within and between trait categories

The comparative study of pollination syndromes suggests that integrated evolution of the many floral traits associated with a given pollination syndrome likely involves coordinated change via a smaller number of genetically-correlated floral modules (Smith, 2016). However, although floral modules have been identified from patterns of genetic correlation and QTL overlap in hybrids in several systems, there is no consensus yet about the prevalence of floral modules within and across trait categories or in different evolutionary contexts (e.g. bee-to-hummingbird vs. outcrosser-selfer or adaptation-with-gene flow vs. allopatric divergence). For example, nectar traits and floral dimensions each form tightly intra-correlated but distinct modules in a transition from outcrossing to selfing in *Ipomaea* (Liao *et al*., 2021)while bee-to-hummingbird transitions run the gamut from highly integrated across all traits in *Mimulus* (Bradshaw *et al*., 1995, 1998; Fishman *et al*., 2013) to largely un-coordinated except by the action of natural selection in the face of gene flow (Wessinger *et al*., 2014, 2023; Kostyun *et al*., 2019). Here, we identify significantly intra-correlated but distinct modules for floral color and dimensions, as well as coordination between the latter module and the reward trait of nectar volume (Fig. 3b). While supporting a role for trait integration and modularity for pollination syndrome evolution, our findings also underline key challenges in applying this conceptual framework at the QTL level.

Elevated QTL overlap within the natural category of pigment traits might suggest pleiotropy as the source of genetic correlation in hybrids as well as trait integration throughout divergent evolution. However, because the carotenoid and anthocyanin/flavanol pigment pathways are biochemically distinct (Grotewold, 2006) and a key multi-pigment supergene has been molecularly dissected in our system, we know that tight integration of pigments traits has causes beyond pleiotropy. In particular, the *YUP-SOLAR-PELAN* supergene on Chr 4 strongly influences all three pigment traits due to tight linkage of adjacent genes (Liang *et al*., 2023). Linkage may also underlie QTL coincidence for the two anthocyanin traits (PLA and NGA) more broadly. MYB transcription factors, including *PELAN* and *SOLAR*, often occur in tandem clusters within plant genomes, providing fertile ground for multiple independent (i.e. linked but potentially non-pleiotropic) mutations affecting anthocyanin production in different tissues. Indeed, such genomic flexibility is key to the proposed importance of both transcription factors (Romani & Moreno, 2021) and gene duplicates (Ohno, 1999) as key loci for evolutionary innovation. Thus, although both pigment and dimensional modules identified in hybrids may reflect pleiotropy, tight linkage is a particularly plausible alternative source for the former. However, their maintenance as modules contributing to pollinator syndrome shifts potentially implicates natural selection acting not only on the individual traits or genes but on trait coordination in the face of gene flow.

Although the tremendous diversity of floral morphologies implies freedom to evolve along many paths, developmental coordination among floral whorls (e.g. petals and stamens) is expected from their serial homology. Indeed, strong genetic correlations (Fig. S2) and elevated QTL overlap among dimensional traits (Figs. 2 & 3) are consistent with a general “flower size” developmental module (Krizek & Anderson, 2013), even if not always due to pleiotropy. Further, coordination of floral dimensions by many multi-trait QTLs is consistent both with parallel interspecific transitions (Fishman *et al*., 2002; Goodwillie *et al*., 2006; Kostyun *et al*., 2019; Liao *et al*., 2021) and with a highly pleiotropic and polygenic basis to standing variation for corolla size traits within *Mimulus* populations (Troth *et al*., 2018). However, the key evolutionary steps in pollination syndrome shifts may often require breaking rather than following the genetic correlations among dimensional traits within populations, and thus may involve rare or novel uncoordinated variants. In particular, the exertion of sexual parts beyond the corolla in hummingbird pollination and loss of stigma-anther distance in autogamous selfers entail changes in the *relative* lengths of different floral whorls, while hummingbird pollination in tubular flowers also involves shifts in corolla length vs. width (Fig. 1). Thus, although many shared loci may influence overall flower size differences during pollination syndrome divergence, key evolutionary shifts in the relative size and position of floral organs must be achieved via specific loci with isolated effects on single traits or via disproportionate shifts in size at many loci.

Thus, despite significant modularity for floral dimensions, the subset of size QTLs with disproportionate effects on different whorls (i.e. less integrated) may be particularly important for divergence in pollination syndromes. For example, the *M. parishii* corolla tube and pistil are the same length, with stamens that are slightly exerted past both (Table 1, Fig. 1), whereas the *M. cardinalis* style extends 13mm past the corolla tube and 3mm past the stamens. This dramatic exertion of the style (a key feature or hummingbird pollination in tubular flowers) implies the action of several PIL-only loci or many multi-trait size loci with slightly greater effects on pistil length (PIL) than stamen (STL) or corolla tube length (CTL). Stigma-anther separation, the key trait for self-pollination, requires similar disproportionality. SAS QTLs exhibit all possible combinations, including joint PIL/STL length QTLs without effects on SAS, regions affecting all three traits, and QTLs that only affect stigma-anther separation (Fig. 2, Table S1). Further, genetic dissection of individual pistil length NILs revealed that one multi-trait QTL (*PIL6&7.2)* could be isolated as a PIL-only factor, while the other (*PIL4.1*) retained parallel effects on multiple length traits. The former is a promising target for dissecting the genetics of mating system evolution *per se*, while the latter is a candidate for overall flower size evolution. More generally, our finding of significant but not particularly high integration of dimensional traits (e.g., QTL overlap indices < ½ those of a similar study in *Ipomoea* outcrosser-selfer hybrids (Liao *et al*., 2021), suggests that genetic coordination of floral dimensions may not be a strong constraint when selection acts on shape as well as size.

In addition to the two modules matching pre-assigned trait categories, our results reveal integration between categories at both the level of the individual QTL (flowering time) and genome-wide (nectar volume). Genome-wide QTL and genetic correlations between flowering time and floral dimensions were not particularly elevated (Fig. 3), but we identified one intriguing genome region that may reflect speed-size co-ordination. FLT4.1, at which *M. parishii* alleles confer much earlier flowering (2*a* = 7.8 to 18.6 days), shares a 0.5-1Mb region on Chr4 with a partially overlapping set of 13 floral-size QTLs across all populations. This co-incidence could be due to linkage among mulitple independent genes, but may reflect a flowering time locus that pleiotropically mediates tradeoffs between speed and size (i.e., fast flowering = small flowers), as in *M. guttatus* (Troth *et al*., 2018). Further genetic dissection of this region promises a clean test of those alternatives. Nectar volume was even more integrated with dimension traits (Fig. 3), with particularly high genetic correlations with flower length traits (Fig. S2), QTL overlap as high as within the dimension module (Fig. 3b) and both NEV QTLs in F_2_s coincident with multi-trait size QTLs (Fig. 2). This result intriguingly contrasts with a recent study of floral integration in *Ipomoea* (Liao *et al*., 2021), where nectar traits formed an evolutionary module only weakly correlated with flower size and few QTL positions were shared. While this difference may in part reflect our measurement of only a single nectar trait, allowing no tests for an even more coordinated nectar-only module, nectar-size integration may reflect the specifics of floral development in tubular flowers. Some components of nectar volume variation (e.g., post-development sugar or water provisioning) may be independent of flower size genes, whereas others (e.g., nectary size) may be directly downstream of developmental shifts causing reduced corollas in *M. parishii*. Further work with additional nectar traits measured in multiple environmental conditions and genetic backgrounds (given non-overlapping RIL and F2 QTLs) will be necessary to tease apart the mechanisms underlying this apparent integration.

### Dissecting the genes underlying polygenic flower size variation in hybrids

To understand the molecular mechanisms of floral evolution and trace their history across species divergence, we must get our hands on the causal variants. NIL generation with phenotypic selection is a common tool for fine-mapping of focal mutants in *Arabidopsis*, and has been used in *Mimulus* to dissect major loci controlling pigment divergence (Yuan *et al*., 2013b, 2016; Liang *et al*., 2022, 2023). Here, pistil length and corolla limb length NILs confirm and refine key dimensional QTLs (see above) and provide a major step towards understanding the molecular basis of poorly understood polygenic traits. This is particularly important for the key mating system trait of pistil length; mutation and hormonal manipulation can alter pistil length dramatically and independently of other floral dimensions (Ding *et al*., 2021), but natural species differences appear highly polygenic and potentially pleiotropic (Tables S1-S2). Although both pistil length NIL intervals still contain many genes, high recombination on chromosome ends in these hybrids (Liang *et al*. 2022; 2023; Sotola *et al*. 2023) makes their further dissection and identification of the causal gene(s) feasible. In contrast, the flower size NIL CLL6&7 (which overlaps with PIL6&7.2) corresponds to a reciprocal translocation (Fishman *et al*., 2013; Stathos & Fishman, 2014; Sotola *et al*., 2023) resistant to further genetic dissection; complementary approaches, such as analyses of gene expression networks conducted on segregating NILs with distinct phenotypes (Langfelder & Horvath, 2008), will be necessary to narrow down functional candidates within this region. Across all traits, additional targeted fine-mapping of even minor QTLs, along with functional approaches, promises a detailed understanding of the many contributors to floral trait divergence in monkeyflowers.

## Conclusions

Overall, our complementary mapping approaches reveal a polygenic genetic architecture even for pollinator attraction (pigment) and reward traits with major leading QTLs, as well as shared QTL hotspots causing strong genetic correlations within pigment and dimensional categories. In addition to enriching understanding of the genetic architecture and modularity of components of pollination syndromes, this work creates a strong foundation for further molecular genetic characterization of floral traits and investigations of the evolutionary genomics of species barriers in this classic model system.

## Acknowledgements

We are grateful to T. Nelson and K. Baesen for assistance with the genotyping and phenotyping at the University of Montana (UM) and to logistical support from the UM ECOR Plant Growth Facility and Genomics Core. We thank C. Liu, M. Opel, and M. Moriarty for plant care in the UConn Botanical Conservatory. We are thankful to S. McCann and P. Beardsley for help in generating the RILs. This work was supported by NSF grants DEB-0846089, DEB-1457763 and OIA-1736249 (to LF), IOS-1827645 (to ALS and YWY), DEB-1350935 (to ALS), and IOS-2319721 (to YWY).

## Author contributions

HC – data collection, analysis, and visualization, drafting of manuscript; CSB – data collection, analysis, and visualization; MS – data collection; VAS – data collection, analysis, and visualization, drafting of initial manuscript; ALS – conceptualization, data collection and interpretation, editing of manuscript; YWY – conceptualization, data collection and interpretation, drafting and editing of manuscript; LF – conceptualization, data collection, analysis, visualization and interpretation, drafting and editing of manuscript. All authors read and are accountable for the submitted manuscript.

## Data availability

The raw sequence data (PRJNA1003462, PRJNA948041) and individual genotypes (doi:10.5061/dryad.v6wwpzh1m) used for generating linkage maps are publicly available. The phenotype matrices used for quantitative genetics and QTL mapping are archived at Dryad https://datadryad.org/stash/share/lDhHN-mqGkS5zdMy9QIcCWuhDNzqm1npxaDtyjAhO6k and will be fully released upon publication. The raw sequencing data used to identify the potential DNA fragments responsible for the long pistils of the PIL NIL and the large flowers of the CLL NIL have been deposited to NCBI under the accession numbers PRJNA1116746 and PRJNA1117077, respectively, and will be released upon publication.

## References

Angert AL. 2009. The niche, limits to species’ distributions, and spatiotemporal variation in demography across the elevation ranges of two monkeyflowers. Proceedings of the National Academy of Sciences of the United States of America 106: 19693–19698.

Angert AL, Schemske DW. 2005. The evolution of species’ distributions: reciprocal transplants across the elevation ranges of Mimulus cardinalis and M. lewisii. Evolution 59: 1671–1684.

Barrett SCH. 2002. The evolution of plant sexual diversity. Nature Reviews Genetics 3: 274–284.

Berardi AE, Esfeld K, Jäggi L, Mandel T, Cannarozzi GM, Kuhlemeier C. 2021. Complex evolution of novel red floral color in Petunia. The Plant Cell 33: 2273–2295.

Bleiweiss R. 2001. Mimicry on the QT(L): genetics of speciation in Mimulus. Evolution 55: 1706–1709.

Bradshaw HD, Otto KG, Frewen BE, McKay JK, Schemske DW. 1998. Quantitative Trait Loci Affecting Differences in Floral Morphology Between Two Species of Monkeyflower (Mimulus). Genetics 149: 367–382.

Bradshaw HD, Schemske DW. 2003. Allele substitution at a flower colour locus produces a pollinator shift in monkeyflowers. Nature 426: 176–178.

Bradshaw HD, Wilbert SM, Otto KG, Schemske DW. 1995. Genetic mapping of floral traits associated with reproductive isolation in monkeyflowers (Mimulus). Nature 376: 762–765.

Byers KJRP, Vela JP, Peng F, Riffell JA, Bradshaw HD. 2014. Floral volatile alleles can contribute to pollinator-mediated reproductive isolation in monkeyflowers (Mimulus). The Plant Journal 80: 1031–1042.

Chen H, Yuan Y-W. 2024. Genetic basis of nectar guide trichome variation between bumblebee- and self-pollinated monkeyflowers (Mimulus): role of the MIXTA-like gene GUIDELESS. BMC Plant Biology 24: 62.

Dellinger AS. 2020. Pollination syndromes in the 21st century: where do we stand and where may we go? New Phytologist 228: 1193–1213.

Ding B, Li J, Gurung V, Lin Q, Sun X, Yuan Y. 2021. The leaf polarity factors SGS3 and YABBYs regulate style elongation through auxin signaling in Mimulus lewisii. New Phytologist 232: 2191–2206.

Ding B, Patterson EL, Holalu SV, Li J, Johnson GA, Stanley LE, Greenlee AB, Peng F, Bradshaw HD, Blinov ML, et al. 2020. Two MYB Proteins in a Self-Organizing Activator-Inhibitor System Produce Spotted Pigmentation Patterns. Current Biology 30: 802–814.

Edwards MB, Choi GPT, Derieg NJ, Min Y, Diana AC, Hodges SA, Mahadevan L, Kramer EM, Ballerini ES. 2021. Genetic architecture of floral traits in bee- and hummingbird-pollinated sister species of Aquilegia (columbine). Evolution 75: 2197–2216.

Fenster CB, Armbruster WS, Wilson P, Dudash MR, Thomson JD. 2004. Pollination syndromes and floral specialization. Annual Review of Ecology, Evolution, and Systematics 35: 375–403.

Fetscher AE. 2001. Resolution of male-female conflict in an hermaphroditic flower. Proceedings of the Royal Society of London. Series B: Biological Sciences 268: 525–529.

Fishman L, Beardsley P, Stathos A, Williams CF, Hill JP. 2015. The genetic architecture of traits associated with the evolution of self-pollination in Mimulus. New Phytologist 205: 907–917.

Fishman L, Kelly AJ, Willis JH. 2002. Minor quantitative trait loci underlie floral traits associated with mating system divergence in Mimulus. Evolution 56: 2138–2155.

Fishman L, McIntosh M, Nelson TC, Baesen K, Finseth FR, Stark-dykema E. 2024. Undoing the ‘nasty: dissecting touch-sensitive stigma movement (thigmonasty) and its loss in self-pollinating monkeyflowers. Biorxiv.

Fishman L, Stathos A, Beardsley P, Williams CF, Hill JP. 2013. Chromosomal rearrangements and the genetics of reproductive barriers in Mimulus (monkeyflowers). Evolution 67: 2547–2560.

Friedman J, Hart KS, Bakker MC den. 2017. Losing one’s touch: Evolution of the touch-sensitive stigma in the Mimulus guttatus species complex. American Journal of Botany 104: 335–341.

Glover BJ, Perez-Rodriguez M, Martin C. 1998. Development of several epidermal cell types can be specified by the same MYB-related plant transcription factor. Development 125: 3497–3508.

Goodwillie C, Kalisz S, Eckert CG. 2005. The evolutionary enigma of mixed mating systems in plants: occurrence, theoretical explanations, and empirical evidence. Annual Review of Ecology, Evolution, and Systematics 36: 47–79.

Goodwillie C, Ritland C, Ritland K. 2006. The genetic basis of floral traits associated with mating system evolution in Leptosiphon (Polemoniaceae): an analysis of quantitative trait loci. Evolution 60: 491–504.

Grotewold E. 2006. The genetics and biochemistry of floral pigments. Annual Review of Plant Biology 57: 761– 780.

Hermann K, Klahre U, Moser M, Sheehan H, Mandel T, Kuhlemeier C. 2013. Tight genetic linkage of prezygotic barrier loci creates a multifunctional speciation island in Petunia. Current Biology 23: 873–877.

Hiesey W, Nobs M, Bjorkman O. 1971. Experimental studies on the nature of species: 5. Biosystematics, genetics and physiological ecology of the Erythranthe section of Mimulus. Carnegie Institution.

Klahre U, Gurba A, Hermann K, Saxenhofer M, Bossolini E, Guerin PM, Kuhlemeier C. 2011. Pollinator choice in Petunia depends on two major genetic Loci for floral scent production. Current Biology 21: 730–739.

Kostyun JL, Gibson MJS, King CM, Moyle LC. 2019. A simple genetic architecture and low constraint allow rapid floral evolution in a diverse and recently radiating plant genus. New Phytologist 223: 1009–1022.

Krizek BA, Anderson JT. 2013. Control of flower size. Journal of Experimental Botany 64: 1427–1437.

Langfelder P, Horvath S. 2008. WGCNA: an R package for weighted correlation network analysis. BMC Bioinformatics 9: 559.

Liang M, Chen W, LaFountain AM, Liu Y, Peng F, Xia R, Bradshaw HD, Yuan Y-W. 2023. Taxon-specific, phased siRNAs underlie a speciation locus in monkeyflowers. Science 379: 576–582.

Liang M, Foster CE, Yuan Y-W. 2022. Lost in translation: Molecular basis of reduced flower coloration in a self-pollinated monkeyflower (Mimulus) species. Science Advances 8: eabo1113.

Liao IT, Rifkin JL, Cao G, Rausher MD. 2021. Modularity and selection of nectar traits in the evolution of the selfing syndrome in Ipomoea lacunosa (Convolvulaceae). New Phytologist 233: 1505–1519.

Lowry DB, Willis JH. 2010. A widespread chromosomal inversion polymorphism contributes to a major life-history transition, local adaptation, and reproductive isolation. PLoS Biology 8: e1000500.

Nelson TC, Muir CD, Stathos AM, Vanderpool DD, Anderson K, Angert AL, Fishman L. 2021a. Quantitative trait locus mapping reveals an independent genetic basis for joint divergence in leaf function, life-history, and floral traits between scarlet monkeyflower (Mimulus cardinalis) populations. American Journal of Botany 108: 844–856.

Nelson TC, Stathos AM, Vanderpool DD, Finseth FR, Yuan Y-W, Fishman L. 2021b. Ancient and recent introgression shape the evolutionary history of pollinator adaptation and speciation in a model monkeyflower radiation (Mimulus section Erythranthe). PLoS Genetics 17: e1009095.

Newcombe FC. 1922. Significance of the Behavior of Sensitive Stigmas. American Journal of Botany 9: 99.

Ohno S. 1999. Gene duplication and the uniqueness of vertebrate genomes circa 1970–1999. Seminars in Cell & Developmental Biology 10: 517–522.

Orr HA. 1998. Testing natural selection vs. genetic drift in phenotypic evolution using quantitative trait locus data. Genetics 149: 2099–2104.

Peng F, Byers KJRP, Bradshaw HD. 2017. Less is more: Independent loss-of-function OCIMENE SYNTHASE alleles parallel pollination syndrome diversification in monkeyflowers (Mimulus). American Journal of Botany 104: 1055–1059.

Ramsey J, Bradshaw HD, Schemske DW. 2003. Components of reproductive isolation between the monkeyflowers Mimulus lewisii and M. cardinalis (Phrymaceae). Evolution 57: 1520–1534.

Rifkin JL, Cao G, Rausher MD. 2021. Genetic architecture of divergence: the selfing syndrome in Ipomoea lacunosa. American Journal of Botany 108: 2038–2054.

Roels SAB, Kelly JK. 2011. Rapid evolution caused by pollinator loss in Mimulus guttatus. Evolution 65: 2541– 2552.

Romani F, Moreno JE. 2021. Molecular mechanisms involved in functional macroevolution of plant transcription factors. New Phytologist 230: 1345–1353.

Schemske DW, Bradshaw HD. 1999. Pollinator preference and the evolution of floral traits in monkeyflowers (Mimulus). Proceedings of the National Academy of Sciences of the United States of America 96: 11910–11915.

Sicard A, Lenhard M. 2011. The selfing syndrome: a model for studying the genetic and evolutionary basis of morphological adaptation in plants. Annals of Botany 107: 1433–1443.

Smith SD. 2016. Pleiotropy and the evolution of floral integration. New Phytologist 209: 80–85.

Sotola VA, Berg CS, Samuli M, Chen H, Mantel SJ, Beardsley PA, Yuan Y-W, Sweigart AL, Fishman L. 2023. Genomic mechanisms and consequences of diverse postzygotic barriers between monkeyflower species. Genetics 225: iyad156.

Stankowski S, Chase MA, McIntosh H, Streisfeld MA. 2023. Integrating top-down and bottom-up approaches to understand the genetic architecture of speciation across a monkeyflower hybrid zone. Molecular Ecology 32: 2041– 2054.

Stathos A, Fishman L. 2014. Chromosomal rearrangements directly cause underdominant F1 pollen sterility in Mimulus lewisii–Mimulus cardinalis hybrids. Evolution 68: 3109–3119.

Stebbins GL. 1970. Adaptive radiation of reproductive characteristics in angiosperms I: Pollination mechanisms. Annual Review of Ecology and Systematics 1: 307–326.

Stuurman J, Hoballah ME, Broger L, Moore J, Basten C, Kuhlemeier C. 2004. Dissection of Floral Pollination Syndromes in Petunia. Genetics 168: 1585–1599.

Troth A, Puzey JR, Kim RS, Willis JH, Kelly JK. 2018. Selective trade-offs maintain alleles underpinning complex trait variation in plants. Science 361: 475–478.

Valle JC del, Gallardo-López A, Buide ML, Whittall JB, Narbona E. 2018. Digital photography provides a fast, reliable, and noninvasive method to estimate anthocyanin pigment concentration in reproductive and vegetative plant tissues. Ecology and Evolution 8: 3064–3076.

Wang S, Basten CJ, Zeng Z-B. 2005. Windows QTL Cartographer 2.5. Dept. of Statistics, North Carolina State Univ.

Wessinger CA, Hileman LC. 2016. Accessibility, constraint, and repetition in adaptive floral evolution. Developmental Biology 419: 175–183.

Wessinger CA, Hileman LC. 2020. Parallelism in Flower Evolution and Development. *Annual Review of Ecology*, Evolution, and Systematics 51: 1–22.

Wessinger CA, Hileman LC, Rausher MD. 2014. Identification of major quantitative trait loci underlying floral pollination syndrome divergence in Penstemon. Philosophical Transactions of the Royal Society B: Biological Sciences 369: 20130349.

Wessinger CA, Katzer AM, Hime PM, Rausher MD, Kelly JK, Hileman LC. 2023. A few essential genetic loci distinguish Penstemon species with flowers adapted to pollination by bees or hummingbirds. PLOS Biology 21: e3002294.

Wessinger CA, Rausher MD. 2012. Lessons from flower colour evolution on targets of selection. Journal of Experimental Botany 63: 5741–5749.

Wilson P, Wolfe AD, Armbruster WS, Thomson JD. 2007. Constrained lability in floral evolution: counting convergent origins of hummingbird pollination in Penstemon and Keckiella. New Phytologist 176: 883–890.

Yeaman S, Whitlock MC. 2011. The genetic architecture of adaptation under migration-selection balance. Evolution 65: 1897–1911.

Yuan Y-W. 2019. Monkeyflowers (Mimulus): new model for plant developmental genetics and evo-devo. New Phytologist 222: 694–700.

Yuan Y-W, Rebocho AB, Sagawa JM, Stanley LE, Bradshaw HD. 2016. Competition between anthocyanin and flavonol biosynthesis produces spatial pattern variation of floral pigments between Mimulus species. Proceedings of the National Academy of Sciences of the United States of America 113: 2448–2453.

Yuan Y-W, Sagawa JM, Frost L, Vela JP, Bradshaw HD. 2014. Transcriptional control of floral anthocyanin pigmentation in monkeyflowers (Mimulus). New Phytologist 204: 1013–1027.

Yuan Y-W, Sagawa JM, Stilio VSD, Bradshaw HD. 2013a. Bulk segregant analysis of an induced floral mutant identifies a MIXTA-like R2R3 MYB controlling nectar guide formation in Mimulus lewisii. Genetics 194: 523–528.

Yuan Y-W, Sagawa JM, Young RC, Christensen BJ, Bradshaw HD. 2013b. Genetic dissection of a major anthocyanin QTL contributing to pollinator-mediated reproductive isolation between sister species of Mimulus. Genetics 194: 255–263.

