## Supporting Information for "The genetic architecture of floral trait divergence between hummingbird- and self-pollinated monkeyflower (*Mimulus*) species"

^*^For correspondence

**The following Supporting Information is available for this article:**

Fig. S1. Phenotypic distributions for traits in the three mapping populations.

Fig. S2. Phenotypic and genetic correlations among traits in F2 hybrids.

Table S1. All QTL effects and locations

Table S2. Phenotypic effects of pistil length NIL

Table S3. Phenotypic effects of corolla limb length NIL

Supplementary Methods S1. UM_F2 and RIL growth conditions

Supplementary Methods S2. UM_F2 and RIL phenotyping

Supplementary Methods S3. Details of NIL generation and genotyping

Fig. S1. Phenotypic distributions for each trait measured in the three *M. parishii* x *M. cardinalis* mapping populations. A) UC_F_2_ grown at the University of Connecticut (n = 253). B) UM_F_2_ grown at the University of Montana (n = 252 total; 249 for dimensional traits and flowering time, 233 for stigma closure, n = 216 for pigment traits) C) RILs grown at the University of Georgia (n = 145 for dimensional traits, 140 for pigment traits, 128 for nectar volume, and 118 for flowering time). Vertical lines indicate the means of the *M. parishii* (pink, dashed), *M. cardinalis* (red, solid) and F_1_ (for UC_F_2_ only; blue, dotted) controls in each population. For the most complete UC_F_2_ growout, where we estimated traditional quantitative genetic metrics, significant deviation from normality by the Shapiro-Wilks test is indicated by asterisks (* < 0.1, *** < 0.001).

A.

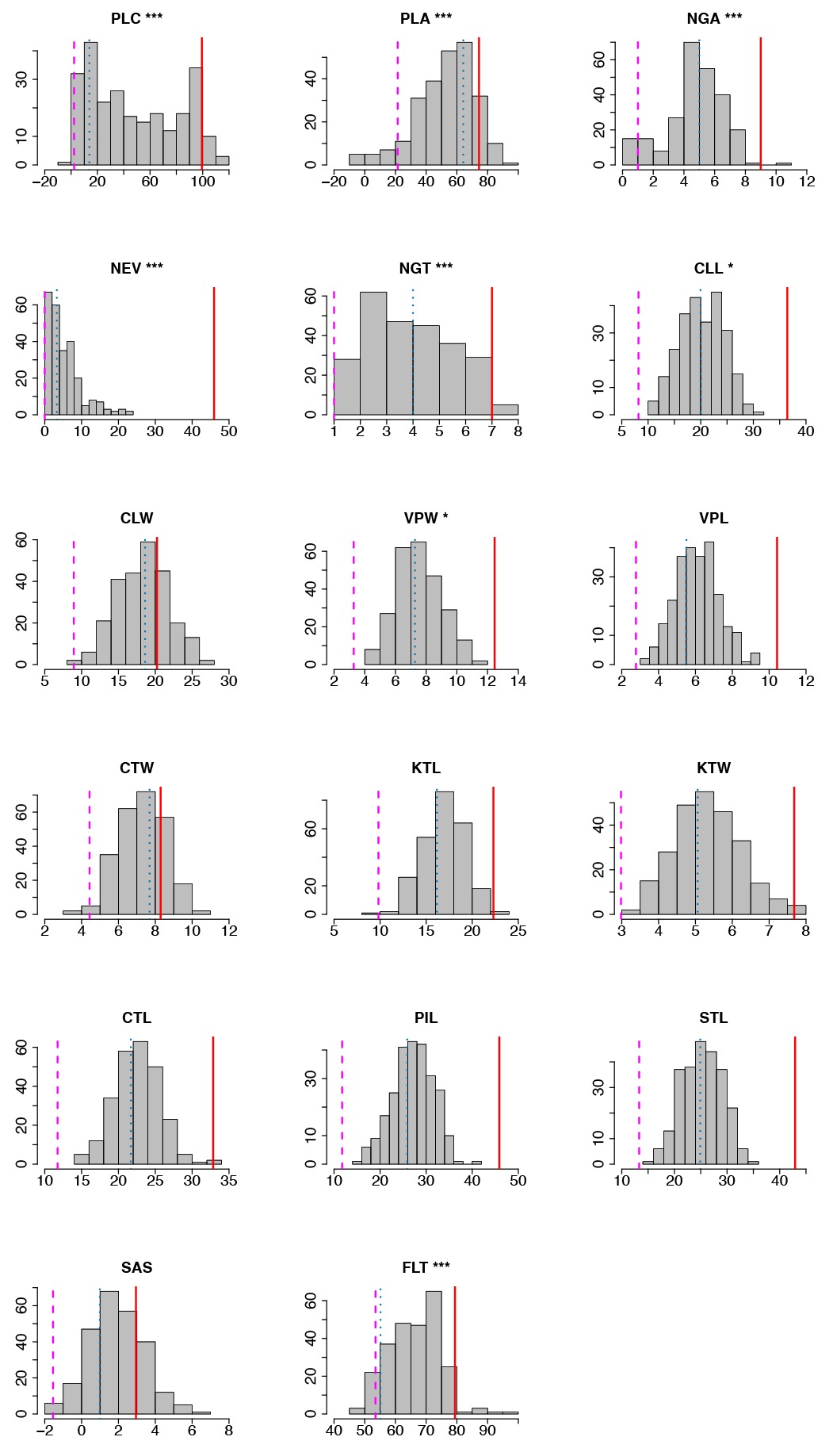

**B.**

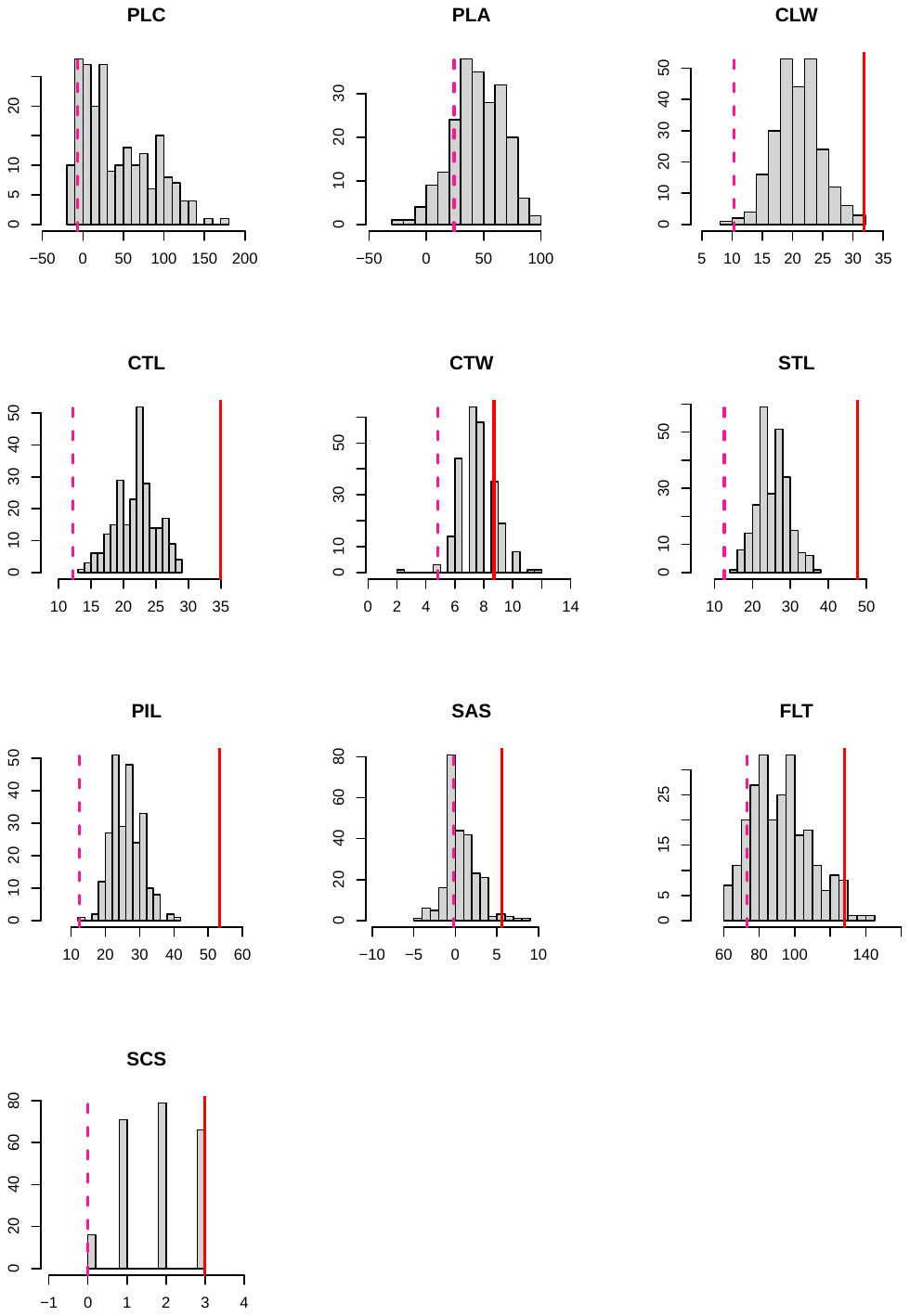

**C.**

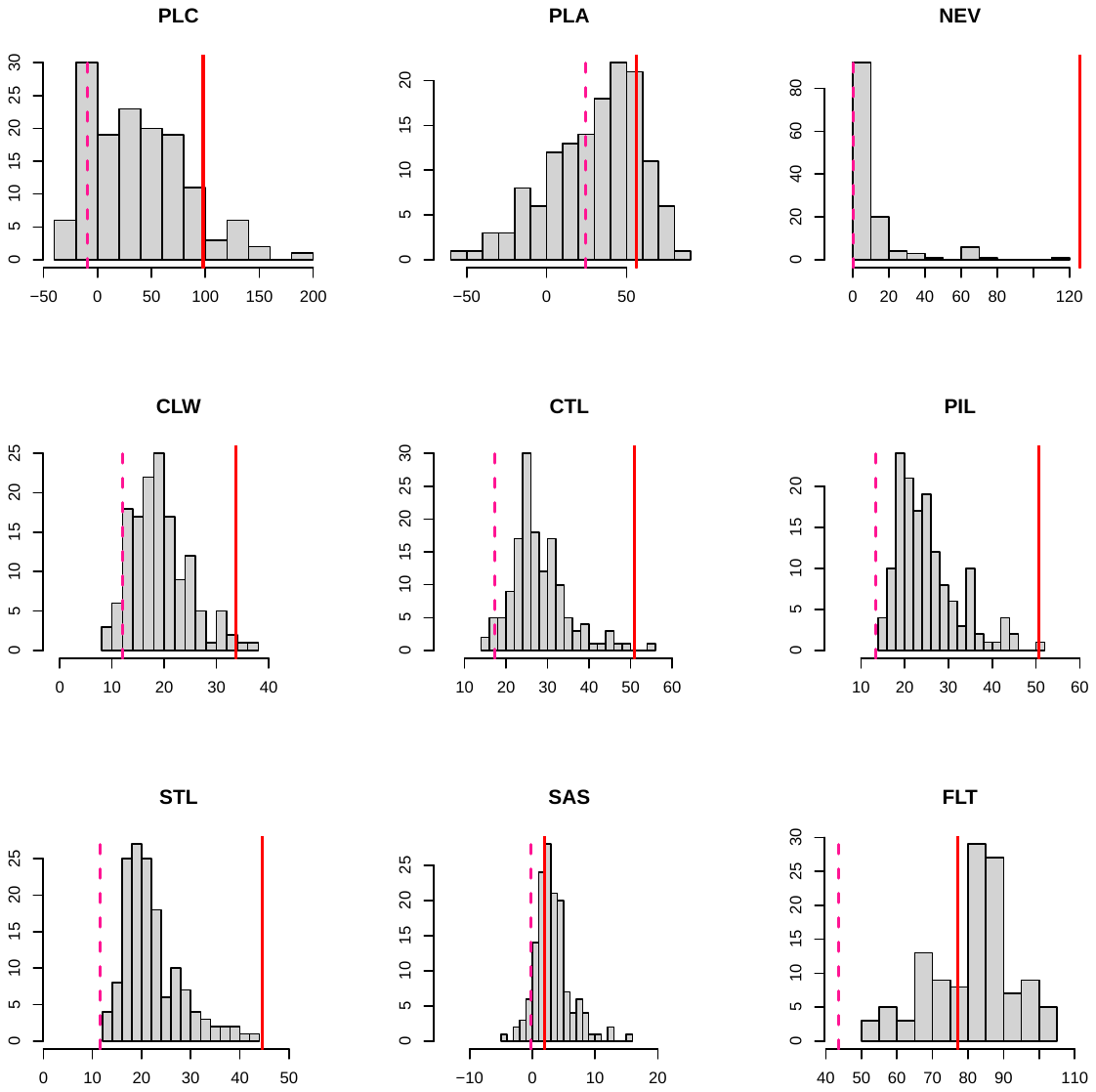

**Fig. S2.** Phenotypic (rP; below the diagonal) and genetic (rG; above the diagonal) correlations among measured traits. PLC: petal lobe carotenoid; PLA: petal lobe anthocyanin; NGA: nectar guide anthocyanin; NEV: nectar volume; NGT: nectar guide trichome; CLL: corolla limb length; CLW: corolla limb width; VPW: ventral petal width; VPL: ventral petal length; CTW: corolla tube width; KTW: calyx tube width; KTL: calyx tube length; CTL: corolla tube length; STL: stamen length; PIL: pistil length; SAS: stigma-anther separation; FLT: flowering time.

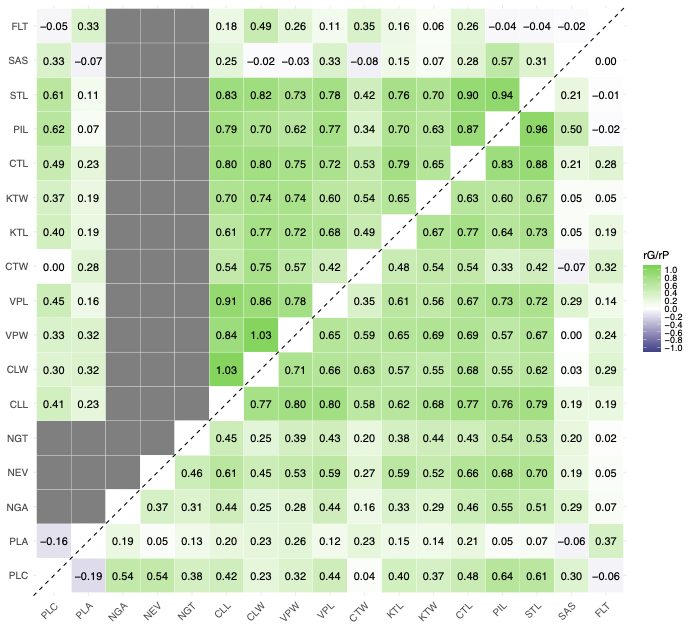

**Table S1.** Floral quantitative trait loci (QTLs) mapped in *Mimulus parishii x* *M. cardinalis* hybrids, organized by trait category as in Fig. 3. **A.** Pigment. **B.** Pollinator reward and handling. **C.** Dimensions, **D.** Flowering time. Columns include the three letter trait codes from Table 1, the mapping population (Pop), the QTL, the additive effect (*a*, positive values indicate alleles move phenotypes toward the larger *M. cardinalis* parent), the dominance effect (*d*), the fraction of the F_2_ variance explained (r^2^), the peak LOD score, the linkage group (LG), the centiMorgan position of QTL LOD peak and its 1.5 LOD-drop confidence interval (cMmin, cMpeak, cMmax) on the corresponding linkage map, Chromosome number (Chrom) and the physical positions of peak and confidence interval on the CE10 *M. cardinalis* v2.0 reference genome (Mbmin, Mbpeak, Mbmax). For QTL comparisons in Fig. 3, the RIL QTL peaks and bounds in cM were translated from RIL linkage map to F_2_ linkage maps positions via their physical bounds. Within shared traits, we binned QTLs across mapping population: ***bold italic*** indicates a QTL was present in all three populations, **bold** indicates a QTL shared by F_2_s, and italics indicate a QTL shared by RILs and one F_2_ population.

**A. Pigment traits**

| Trait | Pop | QTL | *a* | *d* | r^2^ | LOD | LG | cMmin | cMpeak | cMmax | Chrom | Mbmin | Mbpeak | Mbmax |
| --- | --- | --- | --- | --- | --- | --- | --- | --- | --- | --- | --- | --- | --- | --- |
| PLC | UC_F2 | PLC1 | 11.22 | -0.49 | 0.05 | 10.31 | 1 | 18.77 | 32.83 | 34.4 | chr1 | 3.72 | 6.63 | 7.07 |
| PLC | UC_F2 | ***PLC4.1*** | 26.82 | -26.23 | 0.60 | 86.16 | 4 | 28 | 28.11 | 28.22 | chr4 | 7.99 | 8.2 | 8.53 |
| PLC | UC_F2 | **PLC6&7** | 15.01 | -1.23 | 0.06 | 11.79 | 6&7 | 83.04 | 85.89 | 88.34 | chr6 | 55.12 | 56.48 | 57.64 |
| PLC | UM_F2 | ***PLC4.1*** | 30.8 | -32.24 | 0.45 | 43.86 | 4 | 30.2 | 30.67 | 31 | chr4 | 8.53 | 8.9 | 9.16 |
| PLC | UM_F2 | **PLC6&7** | 24.75 | -0.51 | 0.10 | 10.54 | 6&7 | 85.2 | 86.09 | 94.88 | chr6 | 56.19 | 56.65 | 60.55 |
| PLC | RIL | ***PLC4.1*** | 14.51 | -25.69 | 0.12 | 10.29 | 4 | 35.85 | 37.26 | 39.33 | chr4 | 6.72 | 7.98 | 8.6 |
| PLC | RIL | PLC4.2 | -0.71 | -25.27 | 0.05 | 4.34 | 4 | 63.44 | 64.51 | 65.2 | chr4 | 42.34 | 42.62 | 42.75 |
| PLC | RIL | PLC5 | 16.72 | -12.87 | 0.08 | 6.15 | 5 | 79.68 | 84.16 | 86.89 | chr5 | 49.19 | 50.1 | 50.37 |
| PLA | UC_F2 | **PLA3** | 6.86 | 2.48 | 0.06 | 4.71 | 3 | 63.09 | 71.63 | 73.01 | chr3 | 48.26 | 49.36 | 49.55 |
| PLA | UC_F2 | ***PLA4.1*** | 10.82 | 18.13 | 0.14 | 12.06 | 4 | 26.93 | 27.33 | 30.37 | chr4 | 7.65 | 7.99 | 8.69 |
| PLA | UC_F2 | **PLA4.2** | 12.73 | 8.66 | 0.19 | 13.85 | 4 | 70.66 | 71.74 | 71.74 | chr4 | 49.47 | 49.62 | 49.62 |
| PLA | UC_F2 | *PLA6&7* | -13.27 | -6.94 | 0.06 | 4.76 | 6&7 | 89.92 | 91.59 | 91.88 | chr6 | 58.45 | 60.3 | 60.55 |
| PLA | UM_F2 | **PLA3** | 10.79 | -0.36 | 0.11 | 8.14 | 3 | 68.1 | 71.93 | 73.01 | chr3 | 48.96 | 49.43 | 49.55 |
| PLA | UM_F2 | ***PLA4.1*** | 6.61 | 17.76 | 0.13 | 11.44 | 4 | 23.4 | 27.33 | 29.11 | chr4 | 6.58 | 7.99 | 8.2 |
| PLA | UM_F2 | **PLA4.2** | 9.36 | 6.37 | 0.09 | 6.69 | 4 | 66.35 | 66.92 | 71.74 | chr4 | 48.8 | 48.98 | 49.62 |
| PLA | RIL | ***PLA4.1*** | 7.44 | 25.34 | 0.16 | 6.41 | 4 | 29.6 | 33.43 | 36.57 | chr4 | 2.59 | 6.44 | 7.28 |
| PLA | RIL | *PLA6&7* | -15.85 | -4.77 | 0.19 | 6.76 | 6&7 | 50.01 | 50.01 | 50.7 | chr6 | 57.09 | 57.09 | 57.15 |
| NGA | UC_F2 | NGA3 | 1.27 | -0.03 | 0.21 | 25.45 | 3 | 69.37 | 71.63 | 72.13 | chr3 | 49.17 | 49.36 | 49.43 |
| NGA | UC_F2 | NGA4.1 | 2.00 | 0.38 | 0.31 | 34.22 | 4 | 27.61 | 28.11 | 28.61 | chr4 | 7.99 | 8.2 | 8.53 |
| NGA | UC_F2 | NGA5 | 0.73 | 0.56 | 0.04 | 5.74 | 5 | 26.93 | 32.04 | 33.22 | chr5 | 4.76 | 6.65 | 7.2 |
| NGA | UC_F2 | NGA6&7 | 0.62 | -0.19 | 0.05 | 7.79 | 6&7 | 2.37 | 8.46 | 17.21 | chr6 | 0.47 | 1.33 | 2.94 |

**B. Pollinator reward/handling traits**

| Trait | Pop | QTL | *a* | *d* | r^2^ | LOD | LG | cMmin | cMpeak | cMmax | chrom | Mbmin | Mbpeak | Mbmax |
| --- | --- | --- | --- | --- | --- | --- | --- | --- | --- | --- | --- | --- | --- | --- |
| NEV | UC_F2 | NEV1 | 1.14 | 1.43 | 0.05 | 5.31 | 1 | 38.33 | 51.1 | 54.93 | chr1 | 8.61 | 39.10 | 40.14 |
| NEV | UC_F2 | NEV4 | 1.49 | -0.37 | 0.05 | 4.68 | 4 | 50.81 | 53.66 | 60.54 | chr4 | 45.58 | 46.08 | 47.99 |
| NEV | UC_F2 | NEV2 | 2.20 | -0.47 | 0.12 | 11.12 | 2 | 19.18 | 22.62 | 30.38 | chr2 | 4.17 | 4.92 | 6.52 |
| NEV | UC_F2 | NEV6&7 | 3.12 | -1.82 | 0.17 | 16.01 | 6&7 | 155.55 | 157.32 | 158 | chr7 | 15.10 | 16.56 | 17.95 |
| NEV | RIL | NEV5 | 10.90 | -16.31 | 0.21 | 10.97 | 5 | 75.2 | 77.27 | 77.61 | chr5 | 47.48 | 48.36 | 48.84 |
| NEV | RIL | NEV8 | 14.34 | -14.50 | 0.15 | 7.64 | 8 | 37.95 | 46.47 | 50.61 | chr8 | 4.84 | 5.84 | 6.85 |
| NGT | UC_F2 | NGT5 | 0.63 | -0.33 | 0.08 | 7.16 | 5 | 3.74 | 9.93 | 16.42 | chr5 | 0.96 | 1.54 | 2.67 |
| NGT | UC_F2 | NGT6&7.1 | 0.53 | -0.11 | 0.05 | 3.94 | 6&7 | 36.66 | 46.98 | 50.61 | chr6 | 9.99 | 16.95 | 40.74 |
| NGT | UC_F2 | NGT8 | 0.84 | 0.10 | 0.11 | 8.82 | 8 | 53.07 | 55.72 | 59.36 | chr8 | 12.32 | 43.37 | 46.42 |
| NGT | UC_F2 | NGT6&7.2 | 0.53 | 0.25 | 0.04 | 3.86 | 6&7 | 152.51 | 158.79 | 160.95 | chr7 | 12.43 | 19.36 | 33.98 |
| SCS | UM_F2 | SCS5 | 0.30 | 0.23 | 0.06 | 4.10 | 5 | 57.78 | 61.52 | 61.52 | chr5 | 50.92 | 51.63 | 51.63 |
| SCS | UM_F2 | SCS6&7 | 0.31 | 0.27 | 0.05 | 3.80 | 6&7 | 118.02 | 129.03 | 131.19 | chr6 | 47.58 | 48.17 | 52.76 |

**C. Floral dimensions**

|  | Trait | Pop | QTL | | | *a* | *d* | | r^2^ | | LOD | | LG | | cMmin | | cMpeak | | cMmax | | chrom | | Mbmin | | Mbpeak | | Mbmax | |
| --- | --- | --- | --- | --- | --- | --- | --- | --- | --- | --- | --- | --- | --- | --- | --- | --- | --- | --- | --- | --- | --- | --- | --- | --- | --- | --- | --- | --- |
|  | CLL | UC_F2 | CLL1 | | | 1.64 | 0.83 | | 0.06 | | 6.09 | | 1 | | 18.77 | | 30.37 | | 38.33 | | chr1 | | 3.72 | | 6.02 | | 8.61 | |
|  | CLL | UC_F2 | CLL2.1 | | | 1.58 | 0.00 | | 0.07 | | 6.93 | | 2 | | 21.63 | | 25.07 | | 34.9 | | chr2 | | 4.61 | | 5.46 | | 7.86 | |
|  | CLL | UC_F2 | CLL2.2 | | | 1.45 | 0.03 | | 0.06 | | 5.82 | | 2 | | 67.02 | | 69.09 | | 77.73 | | chr2 | | 43.96 | | 45.27 | | 48.52 | |
|  | CLL | UC_F2 | CLL3 | | | 1.73 | -0.33 | | 0.07 | | 7.50 | | 3 | | 39.01 | | 39.51 | | 40.59 | | chr3 | | 12.30 | | 12.98 | | 13.84 | |
|  | CLL | UC_F2 | CLL4 | | | 1.32 | -0.49 | | 0.05 | | 5.55 | | 4 | | 49.86 | | 51.2 | | 71.74 | | chr4 | | 45.43 | | 45.78 | | 49.62 | |
|  | CLL | UC_F2 | CLL5.1 | | | 1.06 | 1.07 | | 0.05 | | 4.69 | | 5 | | 0.01 | | 4.43 | | 16.42 | | chr5 | | 0.34 | | 0.99 | | 2.67 | |
|  | CLL | UC_F2 | CLL5.2 | | | 1.52 | -1.01 | | 0.08 | | 7.38 | | 5 | | 42.55 | | 47.07 | | 48.64 | | chr5 | | 42.69 | | 48.56 | | 49.14 | |
|  | CLL | UC_F2 | CLL6&7.1 | | | 2.09 | 0.02 | | 0.11 | | 9.58 | | 6&7 | | 16.22 | | 17.6 | | 20.94 | | chr6 | | 2.73 | | 3.06 | | 4.26 | |
|  | CLL | UC_F2 | CLL6&7.2 | | | 2.12 | 0.37 | | 0.10 | | 8.96 | | 6&7 | | 146.22 | | 151.23 | | 158.79 | | chr7 | | 5.55 | | 10.67 | | 20.38 | |
|  | CLL | UC_F2 | CLL8 | | | 1.12 | 1.05 | | 0.06 | | 6.05 | | 8 | | 33.42 | | 34.5 | | 43.14 | | chr8 | | 4.86 | | 4.86 | | 7.29 | |
|  | CLW | UC_F2 | **CLW1** | | | 1.10 | 1.09 | | 0.05 | | 5.25 | | 1 | | 20.64 | | 27.32 | | 38.33 | | chr1 | | 3.90 | | 5.17 | | 8.61 | |
|  | CLW | UC_F2 | **CLW2.2** | | | 1.62 | -0.37 | | 0.11 | | 9.29 | | 2 | | 66.04 | | 67.81 | | 71.05 | | chr2 | | 46.77 | | 44.41 | | 46.07 | |
|  | CLW | UC_F2 | *CLW3.2* | | | 1.66 | 0.76 | | 0.10 | | 9.13 | | 3 | | 36.46 | | 39.51 | | 44.71 | | chr3 | | 11.70 | | 12.98 | | 40.12 | |
|  | CLW | UC_F2 | **CLW4** | | | 1.40 | 0.14 | | 0.07 | | 5.95 | | 4 | | 51.2 | | 61.62 | | 66.92 | | chr4 | | 45.78 | | 48.14 | | 48.98 | |
|  | CLW | UC_F2 | ***CLW6&7.1*** | | | 1.53 | 0.40 | | 0.07 | | 6.06 | | 6&7 | | 17.6 | | 19.96 | | 24.38 | | chr6 | | 3.06 | | 3.76 | | 5.48 | |
|  | CLW | UM_F2 | **CLW1** | | | 1.07 | -0.15 | | 0.04 | | 4.84 | | 1 | | 18.78 | | 27.32 | | 36.95 | | chr1 | | 3.72 | | 5.17 | | 8.17 | |
|  | CLW | UM_F2 | CLW2.1 | | | 1.76 | 0.68 | | 0.09 | | 10.65 | | 2 | | 10.33 | | 16.82 | | 23.5 | | chr2 | | 2.35 | | 3.37 | | 5.11 | |
|  | CLW | UM_F2 | **CLW2.2** | | | 1.36 | 0.57 | | 0.05 | | 6.14 | | 2 | | 58.87 | | 62.62 | | 67.81 | | chr2 | | 39.16 | | 41.51 | | 44.41 | |
|  | CLW | UM_F2 | CLW3.1 | | | 1.07 | -0.15 | | 0.03 | | 4.22 | | 3 | | 2.39 | | 8.85 | | 13.67 | | chr3 | | 0.55 | | 2.25 | | 2.96 | |
|  | CLW | UM_F2 | **CLW4** | | | 1.37 | -0.33 | | 0.06 | | 7.60 | | 4 | | 62.91 | | 68.79 | | 71.74 | | chr4 | | 48.28 | | 46.40 | | 49.62 | |
|  | CLW | UM_F2 | ***CLW6&7.1*** | | | 1.32 | 0.01 | | 0.05 | | 6.43 | | 6&7 | | 7.68 | | 12.82 | | 19.96 | | chr6 | | 1.05 | | 2.06 | | 3.76 | |
|  | CLW | UM_F2 | CLW6&7.3 | | | 2.31 | -0.46 | | 0.12 | | 13.83 | | 6&7 | | 90.51 | | 91.59 | | 94.88 | | chr6 | | 59.02 | | 60.30 | | 60.55 | |
|  | CLW | RIL | CLW2.3 | | | 1.12 | -2.89 | | 0.05 | | 4.95 | | 2 | | 75.19 | | 80.02 | | 80.36 | | chr2 | | 43.30 | | 43.70 | | 43.74 | |
|  | CLW | RIL | CLW2.4 | | | 1.14 | -3.03 | | 0.07 | | 6.31 | | 2 | | 84.85 | | 89.67 | | 99.26 | | chr2 | | 44.85 | | 46.29 | | 48.64 | |
|  | CLW | RIL | *CLW3.2* | | | 1.70 | -0.97 | | 0.06 | | 4.98 | | 3 | | 60.68 | | 65.19 | | 70.02 | | chr3 | | 14.50 | | 35.47 | | 41.33 | |
|  | CLW | RIL | ***CLW6&7.1*** | | | 1.99 | -1.72 | | 0.10 | | 7.53 | | 6&7 | | 33.46 | | 36.22 | | 37.6 | | chr6 | | 3.77 | | 4.46 | | 4.88 | |
|  | CLW | RIL | CLW6&7.2 | | | 3.11 | -1.57 | | 0.15 | | 11.37 | | 6&7 | | 63.43 | | 64.5 | | 70.36 | | chr7 | | 10.83 | | 12.82 | | 16.47 | |
|  | Trait | Pop | QTL | | *a* | | *d* | | r^2^ | | LOD | | LG | | cMmin | | cMpeak | | cMmax | | chrom | | Mbmin | | Mbpeak | | Mbmax | |
|  | VPW | UC_F2 | VPW1 | | 0.62 | | 0.08 | | 0.08 | | 9.53 | | 1 | | 30.37 | | 35.77 | | 38.33 | | chr1 | | 6.02 | | 7.75 | | 8.61 | |
|  | VPW | UC_F2 | VPW2 | | 0.93 | | -0.19 | | 0.20 | | 21.71 | | 2 | | 21.93 | | 25.07 | | 28.12 | | chr2 | | 4.76 | | 5.46 | | 6.03 | |
|  | VPW | UC_F2 | VPW4 | | 0.46 | | -0.03 | | 0.04 | | 5.90 | | 4 | | 61.62 | | 61.91 | | 66.92 | | chr4 | | 48.14 | | 48.28 | | 48.98 | |
|  | VPW | UC_F2 | VPW5 | | 0.79 | | -0.24 | | 0.14 | | 15.45 | | 5 | | 44.32 | | 47.07 | | 48.64 | | chr5 | | 47.41 | | 48.56 | | 49.14 | |
|  | VPW | UC_F2 | VPW6&7 | | 0.54 | | 0.12 | | 0.06 | | 7.23 | | 6&7 | | 3.84 | | 8.76 | | 14.16 | | chr6 | | 0.62 | | 1.38 | | 2.39 | |
|  | VPW | UC_F2 | VPW8 | | 0.46 | | 0.31 | | 0.07 | | 8.54 | | 8 | | 33.42 | | 34.5 | | 45.14 | | chr8 | | 4.86 | | 5.05 | | 7.87 | |
|  | VPL | UC_F2 | VPL2.1 | | 0.34 | | 0.11 | | 0.03 | | 3.70 | | 2 | | 2.6 | | 6.73 | | 13.38 | | chr2 | | 0.56 | | 1.82 | | 2.77 | |
|  | VPL | UC_F2 | VPL2.2 | | 0.43 | | 0.08 | | 0.06 | | 6.81 | | 2 | | 63.62 | | 68.89 | | 73.7 | | chr2 | | 42.30 | | 45.01 | | 46.99 | |
|  | VPL | UC_F2 | VPL3 | | 0.41 | | -0.03 | | 0.05 | | 5.53 | | 3 | | 8.26 | | 11.31 | | 17.4 | | chr3 | | 1.81 | | 1.81 | | 3.87 | |
|  | VPL | UC_F2 | VPL4 | | 0.42 | | 0.27 | | 0.04 | | 4.73 | | 4 | | 36.17 | | 37.25 | | 64.27 | | chr4 | | 36.64 | | 38.69 | | 48.54 | |
|  | VPL | UC_F2 | VPL5 | | 0.59 | | -0.23 | | 0.12 | | 13.33 | | 5 | | 28.11 | | 29.58 | | 30.86 | | chr5 | | 5.01 | | 5.72 | | 6.19 | |
|  | VPL | UC_F2 | VPL6&7.1 | | 0.55 | | -0.07 | | 0.10 | | 10.80 | | 6&7 | | 12.98 | | 20.45 | | 22.22 | | chr6 | | 2.06 | | 3.96 | | 4.59 | |
|  | VPL | UC_F2 | VPL6&7.2 | | 0.89 | | -0.02 | | 0.21 | | 21.29 | | 6&7 | | 148.97 | | 149.56 | | 151.43 | | chr7 | | 7.97 | | 9.01 | | 10.84 | |
|  | CTW | UC_F2 | CTW2.2 | | 0.67 | | -0.18 | | 0.14 | | 10.42 | | 2 | | 55.33 | | 56.51 | | 57.98 | | chr2 | | 16.45 | | 16.45 | | 37.18 | |
|  | CTW | UC_F2 | **CTW4** | | 0.54 | | 0.08 | | 0.07 | | 5.85 | | 4 | | 53.66 | | 61.62 | | 71.74 | | chr4 | | 46.08 | | 48.14 | | 49.62 | |
|  | CTW | UC_F2 | *CTW8* | | 0.50 | | 0.12 | | 0.09 | | 6.95 | | 8 | | 0.01 | | 5.51 | | 14.85 | | chr8 | | 0.11 | | 0.84 | | 2.16 | |
|  | CTW | UM_F2 | *CTW2.1* | | 0.58 | | -0.01 | | 0.08 | | 7.17 | | 2 | | 22.62 | | 24.19 | | 29.81 | | chr2 | | 4.92 | | 5.32 | | 6.21 | |
|  | CTW | UM_F2 | CTW2.3 | | 0.46 | | -0.02 | | 0.05 | | 4.06 | | 2 | | 59.56 | | 66.04 | | 69.09 | | chr2 | | 40.23 | | 41.09 | | 45.27 | |
|  | CTW | UM_F2 | **CTW4** | | 0.41 | | -0.07 | | 0.05 | | 4.06 | | 4 | | 54.44 | | 68.79 | | 71.74 | | chr4 | | 46.40 | | 46.40 | | 49.62 | |
|  | CTW | UM_F2 | CTW5 | | -0.54 | | -0.14 | | 0.06 | | 5.07 | | 5 | | 11.72 | | 17.7 | | 24.97 | | chr5 | | 1.80 | | 2.98 | | 4.33 | |
|  | CTW | RIL | *CTW2.1* | | 0.62 | | -0.04 | | 0.11 | | 4.42 | | 2 | | 11.39 | | 15.87 | | 27.56 | | chr2 | | 3.64 | | 4.41 | | 6.41 | |
|  | CTW | RIL | *CTW8* | | 0.64 | | -0.81 | | 0.13 | | 5.08 | | 8 | | 8.63 | | 18.63 | | 27.94 | | chr8 | | 1.57 | | 2.75 | | 3.96 | |
|  | KTW | UC_F2 | KTW2.1 | | 0.42 | | | 0.00 | | 0.09 | | 10.08 | | 2 | | 10.33 | | 19.57 | | 32.05 | | chr2 | | 2.35 | | 4.32 | | 6.95 |
|  | KTW | UC_F2 | KTW2.2 | | 0.35 | | | -0.15 | | 0.07 | | 7.58 | | 2 | | 56.41 | | 57.98 | | 61.62 | | chr2 | | 18.60 | | 37.18 | | 41.51 |
|  | KTW | UC_F2 | KTW4 | | 0.46 | | | 0.00 | | 0.11 | | 11.92 | | 4 | | 57.59 | | 61.62 | | 63.09 | | chr4 | | 47.08 | | 48.14 | | 48.43 |
|  | KTW | UC_F2 | KTW6&7 | | 0.37 | | | 0.08 | | 0.06 | | 6.57 | | 6&7 | | 42.56 | | 46.98 | | 49.53 | | chr6 | | 12.70 | | 16.95 | | 38.54 |
|  | KTW | UC_F2 | KTW8 | | 0.38 | | | 0.10 | | 0.09 | | 10.38 | | 8 | | 23.88 | | 24.08 | | 36.56 | | chr8 | | 3.55 | | 3.67 | | 5.44 |
| Trait | | Pop | QTL | | *a* | | *d* | | r^2^ | | LOD | | LG | | cMmin | | cMpeak | | cMmax | | chrom | | Mbmin | | Mbpeak | | Mbmax | |
| KTL | | UC_F2 | KTL2.1 | | 0.83 | | 0.05 | | 0.05 | | 5.87 | | 2 | | 21.63 | | 23.5 | | 26.94 | | chr2 | | 4.61 | | 5.11 | | 5.86 | |
| KTL | | UC_F2 | KTL2.2 | | 1.22 | | -0.49 | | 0.14 | | 15.23 | | 2 | | 68.89 | | 73.31 | | 73.7 | | chr2 | | 45.01 | | 46.77 | | 46.99 | |
| KTL | | UC_F2 | KTL4 | | 1.55 | | 0.77 | | 0.14 | | 15.13 | | 4 | | 31.45 | | 34.69 | | 36.17 | | chr4 | | 9.16 | | 34.64 | | 36.64 | |
| KTL | | UC_F2 | KTL5 | | 0.78 | | 0.00 | | 0.05 | | 5.78 | | 5 | | 41.37 | | 46.38 | | 53.65 | | chr5 | | 44.51 | | 48.40 | | 50.34 | |
| KTL | | UC_F2 | KTL6&7 | | 0.90 | | 0.48 | | 0.04 | | 4.84 | | 6&7 | | 36.27 | | 46.39 | | 48.55 | | chr6 | | 9.52 | | 15.50 | | 19.42 | |
| KTL | | UC_F2 | KTL8 | | 1.00 | | 0.43 | | 0.11 | | 12.07 | | 8 | | 24.08 | | 28.7 | | 32.83 | | chr8 | | 3.67 | | 4.31 | | 4.31 | |
| CTL | | UC_F2 | CTL1 | | 0.77 | | 0.43 | | 0.03 | | 5.30 | | 1 | | 44.81 | | 46.97 | | 57.98 | | chr1 | | 13.06 | | 35.87 | | 35.87 | |
| CTL | | UC_F2 | **CTL2.1** | | 1.66 | | -0.35 | | 0.14 | | 17.51 | | 2 | | 19.57 | | 22.62 | | 24.19 | | chr2 | | 4.32 | | 4.92 | | 4.61 | |
| CTL | | UC_F2 | CTL3.1 | | 1.92 | | 0.02 | | 0.15 | | 18.76 | | 3 | | 39.51 | | 40.19 | | 42.36 | | chr3 | | 12.98 | | 13.54 | | 20.38 | |
| CTL | | UC_F2 | ***CTL4.1*** | | 1.31 | | 0.22 | | 0.07 | | 9.26 | | 4 | | 6.79 | | 8.36 | | 17.7 | | chr4 | | 1.46 | | 1.81 | | 3.60 | |
| CTL | | UC_F2 | **CTL4.2** | | 1.28 | | -0.13 | | 0.06 | | 8.81 | | 4 | | 46.09 | | 49.86 | | 50.36 | | chr4 | | 43.95 | | 45.43 | | 45.43 | |
| CTL | | UC_F2 | ***CTL5*** | | 1.27 | | -0.08 | | 0.07 | | 9.48 | | 5 | | 44.32 | | 47.27 | | 53.65 | | chr5 | | 47.41 | | 48.56 | | 50.34 | |
| CTL | | UC_F2 | ***CTL6&7*** | | 0.86 | | 0.12 | | 0.03 | | 4.17 | | 6&7 | | 43.44 | | 46.68 | | 48.55 | | chr6 | | 13.43 | | 16.27 | | 19.42 | |
| CTL | | UC_F2 | *CTL8* | | 0.67 | | 0.58 | | 0.04 | | 5.95 | | 8 | | 16.91 | | 30.37 | | 38.62 | | chr8 | | 2.47 | | 4.45 | | 6.28 | |
| CTL | | UM_F2 | **CTL2.1** | | 1.01 | | 0.19 | | 0.04 | | 5.76 | | 2 | | 12.33 | | 16.82 | | 20.57 | | chr2 | | 2.77 | | 3.37 | | 4.46 | |
| CTL | | UM_F2 | CTL2.2 | | 1.01 | | -0.35 | | 0.04 | | 5.80 | | 2 | | 57 | | 67.02 | | 72.53 | | chr2 | | 26.81 | | 43.96 | | 46.53 | |
| CTL | | UM_F2 | CTL3.1 | | 1.42 | | 0.46 | | 0.06 | | 8.67 | | 3 | | 33.61 | | 38.23 | | 40.59 | | chr3 | | 9.29 | | 12.30 | | 13.84 | |
| CTL | | UM_F2 | **CTL4.1b** | 1.32 | | | 0.33 | | 0.04 | | 5.49 | | 4 | | 13.37 | | 23.4 | | 28.11 | | chr4 | | 2.79 | | 6.58 | | 8.20 | |
| CTL | | UM_F2 | **CTL4.2** | 1.19 | | | -0.29 | | 0.06 | | 7.50 | | 4 | | 51.2 | | 54.44 | | 55.44 | | chr4 | | 46.40 | | 46.40 | | 46.85 | |
| CTL | | UM_F2 | ***CTL5*** | -0.81 | | | 0.27 | | 0.03 | | 4.39 | | 5 | | 6.51 | | 10.72 | | 20.45 | | chr5 | | 1.08 | | 1.80 | | 3.40 | |
| CTL | | UM_F2 | ***CTL6&7*** | 1.07 | | | -0.22 | | 0.05 | | 6.82 | | 6&7 | | 119.02 | | 119.59 | | 126.67 | | chr7 | | 0.91 | | 0.99 | | 1.93 | |
| CTL | | RIL | CTL2.2 | 0.83 | | | -4.67 | | 0.08 | | 6.88 | | 2 | | 62.78 | | 64.5 | | 66.91 | | chr2 | | 37.23 | | 38.24 | | 40.94 | |
| CTL | | RIL | CTL3.2 | 1.93 | | | -0.76 | | 0.05 | | 4.12 | | 3 | | 62.78 | | 64.85 | | 64.85 | | chr3 | | 31.69 | | 35.69 | | 36.01 | |
| CTL | | RIL | CTL3.3 | 1.99 | | | -1.44 | | 0.06 | | 5.07 | | 3 | | 69.68 | | 71.06 | | 72.43 | | chr3 | | 40.97 | | 41.52 | | 42.05 | |
| CTL | | RIL | **CTL4.1** | 1.30 | | | -2.73 | | 0.05 | | 4.28 | | 4 | | 14.84 | | 15.53 | | 15.53 | | chr4 | | 3.20 | | 3.16 | | 3.19 | |
| CTL | | RIL | *CTL4.1a* | 0.97 | | | -3.81 | | 0.06 | | 4.78 | | 4 | | 3.77 | | 7.94 | | 9.32 | | chr4 | | 1.53 | | 1.96 | | 2.44 | |
| CTL | | RIL | ***CTL5*** | 2.81 | | | -3.04 | | 0.08 | | 6.41 | | 5 | | 75.89 | | 80.02 | | 85.2 | | chr5 | | 47.81 | | 49.45 | | 50.28 | |
| CTL | | RIL | ***CTL6&7*** | 3.28 | | | -1.60 | | 0.11 | | 7.95 | | 6&7 | | 63.43 | | 64.84 | | 70.36 | | chr7 | | 10.83 | | 11.63 | | 16.47 | |
| CTL | | RIL | *CTL8* | 3.69 | | | -1.67 | | 0.10 | | 6.73 | | 8 | | 42.09 | | 45.47 | | 47.61 | | chr8 | | 5.12 | | 5.84 | | 6.85 | |
| Trait | | Pop | QTL | | *a* | | *d* | | r^2^ | | LOD | | LG | | cMmin | | cMpeak | | cMmax | | chrom | | Mbmin | | Mbpeak | | Mbmax | |
| STL | | UC_F2 | STL1.1 | | 1.24 | | -0.01 | | 0.04 | | 6.88 | | 1 | | 24.18 | | 25.75 | | 36.95 | | chr1 | | 4.42 | | 4.60 | | 8.17 | |
| STL | | UC_F2 | **STL1.2** | | 1.05 | | 0.30 | | 0.04 | | 5.48 | | 1 | | 42.55 | | 51.1 | | 54.93 | | chr1 | | 11.08 | | 39.10 | | 40.14 | |
| STL | | UC_F2 | STL2.1 | | 1.23 | | -0.41 | | 0.05 | | 8.36 | | 2 | | 21.63 | | 23.5 | | 25.37 | | chr2 | | 4.61 | | 5.11 | | 5.61 | |
| STL | | UC_F2 | **STL2.2** | | 1.20 | | 0.09 | | 0.04 | | 6.44 | | 2 | | 42.46 | | 50.52 | | 57 | | chr2 | | 10.50 | | 10.50 | | 34.41 | |
| STL | | UC_F2 | STL2.3 | | 1.22 | | -0.50 | | 0.05 | | 8.03 | | 2 | | 79.21 | | 81.17 | | 85 | | chr2 | | 48.69 | | 49.13 | | 50.02 | |
| STL | | UC_F2 | **STL3** | | 1.40 | | -0.31 | | 0.05 | | 7.26 | | 3 | | 36.46 | | 39.51 | | 48.84 | | chr3 | | 11.70 | | 12.98 | | 43.13 | |
| STL | | UC_F2 | STL4.2 | | 2.43 | | 0.59 | | 0.12 | | 15.42 | | 4 | | 28.11 | | 30.11 | | 35.28 | | chr4 | | 8.20 | | 8.53 | | 21.83 | |
| STL | | UC_F2 | *STL5* | | 1.16 | | -0.05 | | 0.04 | | 5.66 | | 5 | | 44.32 | | 47.27 | | 61.52 | | chr5 | | 47.41 | | 48.56 | | 51.48 | |
| STL | | UC_F2 | STL6&7.2 | | 2.05 | | 0.87 | | 0.08 | | 10.64 | | 6&7 | | 46.19 | | 46.39 | | 48.06 | | chr6 | | 14.90 | | 16.27 | | 18.73 | |
| STL | | UC_F2 | *STL6&7.5* | | 1.46 | | 0.41 | | 0.05 | | 7.20 | | 6&7 | | 148.97 | | 151.23 | | 160.17 | | chr7 | | 7.97 | | 10.67 | | 24.99 | |
| STL | | UC_F2 | **STL8.1** | | 1.93 | | 0.74 | | 0.14 | | 17.36 | | 8 | | 33.42 | | 34.5 | | 35 | | chr8 | | 4.86 | | 5.05 | | 5.21 | |
| STL | | UM_F2 | **STL1.2** | | 1.17 | | -0.17 | | 0.04 | | 5.95 | | 1 | | 42.55 | | 49.23 | | 59.98 | | chr1 | | 11.08 | | 37.76 | | 41.67 | |
| STL | | UM_F2 | **STL2.2** | | 0.95 | | 0.37 | | 0.02 | | 3.66 | | 2 | | 52.09 | | 61.44 | | 70.17 | | chr2 | | 13.38 | | 41.51 | | 45.77 | |
| STL | | UM_F2 | **STL3** | | 1.16 | | 0.97 | | 0.04 | | 6.42 | | 3 | | 33.61 | | 38.23 | | 44.03 | | chr3 | | 9.29 | | 12.30 | | 39.63 | |
| STL | | UM_F2 | STL4.1 | | 1.76 | | 0.00 | | 0.06 | | 9.07 | | 4 | | 9.74 | | 14.37 | | 15.93 | | chr4 | | 1.56 | | 3.03 | | 3.26 | |
| STL | | UM_F2 | STL4.3 | | 1.03 | | 0.05 | | 0.02 | | 3.94 | | 4 | | 42.95 | | 54.44 | | 57.79 | | chr4 | | 42.97 | | 46.40 | | 46.40 | |
| STL | | UM_F2 | STL6&7.3 | | 2.11 | | 0.54 | | 0.07 | | 9.51 | | 6&7 | | 57.88 | | 61.44 | | 61.64 | | chr6 | | 45.49 | | 45.96 | | 46.65 | |
| STL | | UM_F2 | STL6&7.6 | 1.20 | | | -0.25 | | 0.03 | | 5.57 | | 6&7 | | 121.18 | | 123.52 | | 128.34 | | chr7 | | 1.10 | | 1.51 | | 2.11 | |
| STL | | UM_F2 | **STL8.1** | 1.81 | | | -0.09 | | 0.07 | | 10.43 | | 8 | | 34.42 | | 38.62 | | 42.39 | | chr8 | | 4.86 | | 6.28 | | 7.29 | |
| STL | | RIL | STL2.4 | 1.46 | | | -2.73 | | 0.08 | | 7.43 | | 2 | | 85.85 | | 87.26 | | 90.71 | | chr2 | | 44.85 | | 45.70 | | 46.93 | |
| STL | | RIL | *STL5* | 1.63 | | | -2.20 | | 0.05 | | 4.78 | | 5 | | 77.61 | | 84.16 | | 89.34 | | chr5 | | 48.67 | | 50.10 | | 50.65 | |
| STL | | RIL | STL6&7.1 | 1.93 | | | -1.76 | | 0.10 | | 8.47 | | 6&7 | | 31.7 | | 35.88 | | 36.57 | | chr6 | | 3.86 | | 4.40 | | 4.79 | |
| STL | | RIL | STL6&7.4 | 3.58 | | | -0.53 | | 0.12 | | 11.81 | | 6&7 | | 64.15 | | 64.84 | | 67.6 | | chr7 | | 11.90 | | 11.63 | | 15.42 | |
| STL | | RIL | *STL6&7.5* | 3.51 | | | -0.48 | | 0.15 | | 12.24 | | 6&7 | | 69.29 | | 70.01 | | 70.36 | | chr7 | | 15.42 | | 15.97 | | 16.47 | |
| STL | | RIL | STL8.2 | 2.34 | | | -1.74 | | 0.08 | | 6.38 | | 8 | | 56.59 | | 72.07 | | 72.07 | | chr8 | | 10.43 | | 46.85 | | 46.92 | |

| Trait | Pop | QTL | | *a* | *d* | r^2^ | LOD | LG | cMmin | cMpeak | cMmax | chrom | Mbmin | Mbpeak | Mbmax |
| --- | --- | --- | --- | --- | --- | --- | --- | --- | --- | --- | --- | --- | --- | --- | --- |
| PIL | UC_F2 | PIL1 | | 0.92 | 0.68 | 0.02 | 4.11 | 1 | 22.61 | 24.18 | 60.53 | chr1 | 4.05 | 4.42 | 42.31 |
| PIL | UC_F2 | PIL2.1 | | 0.76 | -0.52 | 0.02 | 4.03 | 2 | 19.18 | 22.62 | 32.44 | chr2 | 4.17 | 4.92 | 7.16 |
| PIL | UC_F2 | **PIL2.2** | | 1.09 | -0.30 | 0.03 | 6.14 | 2 | 68.89 | 81.17 | 87.66 | chr2 | 45.01 | 49.13 | 51.16 |
| PIL | UC_F2 | **PIL3** | | 2.10 | 0.62 | 0.09 | 16.24 | 3 | 36.46 | 39.51 | 40.59 | chr3 | 11.70 | 12.98 | 13.84 |
| PIL | UC_F2 | PIL4.1a | | 2.84 | 0.19 | 0.14 | 22.28 | 4 | 15.93 | 17.7 | 19.56 | chr4 | 3.26 | 3.60 | 3.94 |
| PIL | UC_F2 | **PIL4.4** | | 1.40 | -0.10 | 0.04 | 7.49 | 4 | 47.86 | 52.18 | 61.62 | chr4 | 44.51 | 45.93 | 48.14 |
| PIL | UC_F2 | *PIL5* | | 1.55 | -0.31 | 0.05 | 11.76 | 5 | 44.32 | 47.07 | 47.27 | chr5 | 47.41 | 48.56 | 48.56 |
| PIL | UC_F2 | PIL6&7.1 | | 2.15 | 0.28 | 0.08 | 13.75 | 6&7 | 45.8 | 46.39 | 47.57 | chr6 | 14.73 | 15.50 | 17.85 |
| PIL | UC_F2 | PIL6&7.5 | | 2.12 | 0.30 | 0.08 | 13.34 | 6&7 | 89.23 | 90.51 | 91.88 | chr6 | 57.92 | 59.02 | 60.55 |
| PIL | UC_F2 | ***PIL8*** | | 2.10 | 0.66 | 0.12 | 19.83 | 8 | 30.37 | 31.94 | 35.67 | chr8 | 4.45 | 4.66 | 5.21 |
| PIL | UM_F2 | **PIL2.2** | | 0.99 | 0.53 | 0.02 | 4.80 | 2 | 78.73 | 87.48 | 89.03 | chr2 | 48.69 | 50.69 | 51.37 |
| PIL | UM_F2 | **PIL3** | | 1.99 | 0.75 | 0.07 | 13.54 | 3 | 35.4 | 40.2 | 43.24 | chr3 | 10.12 | 13.54 | 38.44 |
| PIL | UM_F2 | PIL4.1b | | 2.55 | -0.05 | 0.12 | 21.38 | 4 | 10.74 | 12.9 | 14.95 | chr4 | 2.50 | 2.79 | 3.03 |
| PIL | UM_F2 | **PIL4.4** | | 1.46 | -0.40 | 0.05 | 9.70 | 4 | 53.66 | 54.44 | 59.47 | chr4 | 46.08 | 46.40 | 47.66 |
| PIL | UM_F2 | PIL6&7.2 | | 1.37 | -0.27 | 0.03 | 5.84 | 6&7 | 27.64 | 34.89 | 44.42 | chr6 | 6.28 | 8.90 | 13.83 |
| PIL | UM_F2 | PIL6&7.3 | | 1.48 | 0.42 | 0.02 | 5.06 | 6&7 | 64.66 | 69.77 | 70.77 | chr6 | 48.17 | 49.57 | 49.99 |
| PIL | UM_F2 | PIL6&7.5 | | 1.95 | 0.22 | 0.04 | 7.61 | 6&7 | 148.58 | 149.07 | 152.01 | chr7 | 6.73 | 61.42 | 11.55 |
| PIL | UM_F2 | ***PIL8*** | 1.39 | | 0.51 | 0.05 | 9.49 | 8 | 27.82 | 40.39 | 42.39 | chr8 | 4.04 | 6.69 | 7.29 |
| PIL | RIL | PIL4.1c | 2.22 | | -2.34 | 0.09 | 9.61 | 4 | 2.04 | 5.87 | 9.32 | chr4 | 1.43 | 1.68 | 2.44 |
| PIL | RIL | *PIL5* | 3.00 | | -3.14 | 0.08 | 8.91 | 5 | 78.99 | 80.71 | 83.47 | chr5 | 48.84 | 49.90 | 50.10 |
| PIL | RIL | PIL6&7.4 | 2.14 | | -0.28 | 0.07 | 7.14 | 6&7 | 69.3 | 74.5 | 78.3 | chr6 | 16.82 | 38.95 | 43.06 |
| PIL | RIL | ***PIL8*** | 3.82 | | -1.72 | 0.10 | 9.43 | 8 | 39.98 | 45.47 | 47.61 | chr8 | 4.80 | 5.84 | 6.85 |

| Trait | Pop | QTL | *a* | *d* | r^2^ | LOD | LG | cMmin | cMpeak | cMmax | chrom | Mbmin | Mbpeak | Mbmax |
| --- | --- | --- | --- | --- | --- | --- | --- | --- | --- | --- | --- | --- | --- | --- |
| SAS | UC_F2 | **SAS2.1** | -0.59 | -0.09 | 0.07 | 6.67 | 2 | 10.27 | 14.46 | 19.18 | chr2 | 2.35 | 2.92 | 4.17 |
| SAS | UC_F2 | SAS2.2 | -0.17 | 0.62 | 0.05 | 5.40 | 2 | 45.8 | 56.41 | 57.98 | chr2 | 11.12 | 11.12 | 37.18 |
| SAS | UC_F2 | *SAS3.1* | 0.61 | 0.08 | 0.07 | 6.46 | 3 | 11.31 | 19.17 | 21.43 | chr3 | 2.56 | 4.63 | 4.63 |
| SAS | UC_F2 | *SAS4.2* | 0.88 | 0.32 | 0.14 | 12.38 | 4 | 8.76 | 9.74 | 10.91 | chr4 | 1.95 | 2.25 | 2.50 |
| SAS | UC_F2 | SAS6&7.1 | 0.52 | 0.05 | 0.04 | 3.88 | 6&7 | 39.74 | 42.56 | 51.69 | chr6 | 10.61 | 12.70 | 42.59 |
| SAS | UC_F2 | SAS6&7.3 | 0.82 | -0.58 | 0.12 | 11.25 | 6&7 | 154.08 | 158.11 | 160.95 | chr7 | 13.80 | 17.95 | 33.98 |
| SAS | UM_F2 | SAS1 | -0.66 | -0.05 | 0.06 | 4.54 | 1 | 43.44 | 49.23 | 52.08 | chr1 | 12.12 | 37.76 | 39.59 |
| SAS | UM_F2 | **SAS2.1** | -0.66 | 0.11 | 0.05 | 4.55 | 2 | 1.6 | 4.73 | 16.23 | chr2 | 0.35 | 0.78 | 3.30 |
| SAS | UM_F2 | SAS3.2 | 0.73 | -0.26 | 0.04 | 3.71 | 3 | 38.23 | 40.2 | 44.03 | chr3 | 12.30 | 13.54 | 39.63 |
| SAS | UM_F2 | SAS4.1 | 0.81 | 0.10 | 0.07 | 6.02 | 4 | 0.01 | 3.84 | 7.48 | chr4 | 0.42 | 1.04 | 1.56 |
| SAS | UM_F2 | SAS6&7.2 | 0.87 | 0.06 | 0.07 | 5.67 | 6&7 | 56.02 | 60.44 | 65.66 | chr6 | 44.79 | 45.96 | 48.17 |
| SAS | RIL | *SAS3.1* | 1.07 | -0.73 | 0.12 | 5.52 | 3 | 16.18 | 21.39 | 22.77 | chr3 | 3.08 | 3.59 | 3.98 |
| SAS | RIL | *SAS4.2* | 0.93 | -1.85 | 0.12 | 5.63 | 4 | 6.56 | 19.32 | 20.01 | chr4 | 1.90 | 3.77 | 3.97 |
| SAS | RIL | SAS4.3 | 1.19 | 1.46 | 0.15 | 6.91 | 4 | 84.87 | 87.21 | 90.05 | chr4 | 46.76 | 46.81 | 47.34 |
| SAS | RIL | SAS5 | 1.37 | -0.66 | 0.10 | 4.79 | 5 | 78.99 | 80.02 | 83.47 | chr5 | 48.84 | 49.45 | 50.10 |

**D. Flowering time**

| Trait | Pop | QTL | | a | d | r2 | LOD | LG | cMmin | cMpeak | cMmax | chrom | Mbmin | Mbpeak | Mbmax |
| --- | --- | --- | --- | --- | --- | --- | --- | --- | --- | --- | --- | --- | --- | --- | --- |
| FLT | UC_F2 | FLT2.3 | | 3.85 | -1.01 | 0.11 | 10.27 | 2 | 63.62 | 65.26 | 73.31 | chr2 | 42.30 | 43.28 | 46.77 |
| FLT | UC_F2 | FLT3.2 | | 3.09 | -1.05 | 0.07 | 7.11 | 3 | 63.09 | 68.69 | 69.87 | chr3 | 48.26 | 49.08 | 49.26 |
| FLT | UC_F2 | ***FLT4.1*** | 4.81 | | 1.37 | 0.14 | 12.41 | 4 | 69.2 | 70.66 | 71.74 | chr4 | 49.29 | 49.47 | 49.62 |
| FLT | UC_F2 | FLT6&7.1 | -3.89 | | -1.98 | 0.11 | 10.24 | 6&7 | 129.52 | 132.17 | 132.76 | chr7 | 2.35 | 2.70 | 2.83 |
| FLT | UC_F2 | *FLT8* | -4.32 | | -1.29 | 0.14 | 12.59 | 8 | 38.62 | 49.53 | 53.07 | chr8 | 6.28 | 10.12 | 12.50 |
| FLT | UM_F2 | FLT2.1 | 6.60 | | 1.60 | 0.07 | 5.80 | 2 | 31.38 | 34.9 | 39.52 | chr2 | 6.71 | 7.86 | 9.49 |
| FLT | UM_F2 | FLT3.1 | 6.22 | | -5.86 | 0.09 | 7.81 | 3 | 50.02 | 55.62 | 58.27 | chr3 | 43.34 | 46.55 | 47.20 |
| FLT | UM_F2 | ***FLT4.1*** | 9.68 | | -0.44 | 0.15 | 12.34 | 4 | 65.27 | 71.74 | 71.74 | chr4 | 48.54 | 49.62 | 49.62 |
| FLT | UM_F2 | FLT6&7.2 | -6.88 | | -6.39 | 0.11 | 9.31 | 6&7 | 184.45 | 184.55 | 189.75 | chr7 | 43.09 | 43.36 | 43.09 |
| FLT | RIL | FLT2.2 | 4.99 | | -5.74 | 0.17 | 6.35 | 2 | 50.71 | 57.26 | 62.09 | chr2 | 13.25 | 19.95 | 37.23 |
| FLT | RIL | ***FLT4.1*** | 3.91 | | -6.62 | 0.12 | 4.35 | 4 | 106.53 | 109.02 | 110.4 | chr4 | 49.21 | 49.38 | 49.61 |
| FLT | RIL | *FLT8* | -4.76 | | -5.33 | 0.12 | 4.35 | 8 | 55.87 | 62.11 | 67.28 | chr8 | 9.92 | 39.51 | 44.98 |

**Table S2 Phenotypic statistics (mm, mean ± SD) of 9 kinds of genotypes generated by *PIL4.1* (on Chr4) and *PIL6&7.2* (on Chr6) loci (n = 10 mature flowers for each genotype).** C: homozygous for *M. cardinalis*; H: heterozygous; P: homozygous for *M. parishii*; For genotypes, the *PIL 4.1* genotypes were shown first, followed by the *PIL 6&7.2* genotypes. CLL: corolla limb length; CTL: corolla tube length; PIL: pistil length; STL: stamen length. SAS: stigma-anther separation.

| Genotypes | CLL (mm) | CTL (mm) | PIL (mm) | STL (mm) | SAS (mm) |
| --- | --- | --- | --- | --- | --- |
| *M. parishii* | 13.995±1.28 | 15.545±1.17 | 15.272±0.75 | 15.991±0.90 | -0.719±0.34 |
| C; C | 14.264±0.84 | 17.017±0.94 | 19.148±0.68 | 17.577±0.56 | 1.571±0.32 |
| C; P | 13.158±0.85 | 17.001±0.64 | 18.249±0.66 | 17.181±0.51 | 1.068±0.27 |
| C; H | 14.181±0.35 | 17.349±0.62 | 18.839±0.60 | 17.567±0.49 | 1.272±0.35 |
| H; C | 14.715±0.96 | 16.558±0.80 | 17.983±0.93 | 17.111±0.82 | 0.872±0.32 |
| H; P | 14.089±0.73 | 16.789±0.76 | 16.893±0.55 | 16.656±0.59 | 0.237±0.19 |
| H; H | 14.552±0.97 | 16.336±0.90 | 17.507±0.65 | 16.84±0.69 | 0.667±0.26 |
| P; C | 14.449±0.98 | 14.839±0.59 | 16.122±0.44 | 15.482±0.50 | 0.64±0.20 |
| P; P | 13.232±1.43 | 15.385±0.75 | 15.238±0.61 | 15.825±0.53 | -0.587±0.24 |
| P; H | 13.907±1.17 | 15.893±1.05 | 15.905±0.92 | 15.824±0.84 | 0.081±0.24 |

**Table S3 Phenotypic statistics (mean ± SD) of the *CLL6&7.2* NIL (n = 10 mature flowers for each genotype).** CTL: corolla tube length; PIL: pistil length; STL: stamen length; SAS: stigma-anther separation; CLL: corolla limb length.

| Genotypes | CTL (mm) | PIL (mm) | STL (mm) | SAS (mm) | CLL (mm) |
| --- | --- | --- | --- | --- | --- |
| *M. parishii* | 13.274±0.31 | 13.111±0.35 | 13.628±0.37 | -0.517±0.20 | 11.075±0.46 |
| *CLL6&7.2* NIL | 15.259±0.61 | 15.602±0.42 | 15.376±0.64 | 0.226±0.25 | 12.682±0.40 |

**Methods S1**

**Plant growth conditions for QTL mapping -** F_2_ hybrids were grown in two separate greenhouse common gardens at the University of Connecticut (UC) and the University of Montana (UM). At UC, seeds (CE10, PAR, F_1_ hybrids, F_2_ hybrids, and NILs) were sown into 98 cell trays (1.25"sq x 2"H), and then after four weeks, all seedlings were transferred to 18 pocket trays (3.25" sq x 3.14"H) and grown to flowering. At UM, F_2_ seeds (and a few CE10 and PAR controls) were germinated on wet sand in Petri dishes, seedlings transplanted into 3” square pots filled with Sunshine #4 soil-less potting mix, and plant grown to first flower and beyond (life history and plant architecture traits presented elsewhere). The RIL mapping population was grown at the University of Georgia (UGA) with seeds sown into 2.5” pots filled with Metromix 830 soil. Seedlings were transplanted into 4” pots and grown to flowering. In all three greenhouses, plants were grown under natural light supplemented with sodium lamps to a 16‐h daylength and fertilized regularly (UC: 2-3 times per week, UM and UGA: weekly).

**Methods S2**

**Phenotypic measurements for QTL mapping: UM_F_2_s and RILs -** In addition to pollen number and viability (Sotola *et al.*, 2023) and life history traits to be presented elsewhere, the UM_F_2_ grow-out measured 13 traits shared with the UC_F_2_ growout: flowering time, two floral pigment traits (PLA and PLC), and six floral dimension traits (corolla length and width, corolla tube width, and pistil and stamen length) plus stigma-anther separation. PLA and PLC were measured and calculated as described above, except that we scanned one of the two lateral petals. The corolla dimension traits were measured with an engineering ruler on one flower of the first pair on the day it opened. Stigma closure is described in the main text.

In the RIL growout at UGA (n = 145 RILs, 18 each of CE10 and PAR parents and F_1_ hybrids), we recorded day to first flower (flowering time) and measured two floral pigment traits (PLA and PLC), nectar volume, and six floral dimension traits (corolla length and width, corolla tube width, and pistil and stamen length, stigma-anther separation.) We measured floral pigment traits on the ventral petal as described above and used an engineering ruler to measure the floral dimensional traits on one flower from the first pair. To estimate nectar volume, we used a capillary tube to collect nectar from two flowers on their first day of opening and took the average; nectar was measured between 1:00 PM - 3:00 PM after watering in the morning at 8:00 AM - 9:00 AM. Other than pistil length (PIL), which includes the stigma lobes in the UC_F_2_ and UGA_RIL measurements but not in the UM_F_2_ dataset, and thus stigma-anther separation (SAS), traits with a shared abbreviation represent the same floral dimension.

**Methods S3**

**Construction and characterization of NILs** -To construct pistil length NILs in the *M. parishii* genetic background, we chose an (ungenotyped) F_2_ individual with overall similarity to *M. parishii* in both floral and vegetative traits, but with conspicuously longer pistil, for serial backcrossing to *M. parishii* (as maternal parent). From each backcross growout of ~95 plants, we selected one individual for the next round that closely resembled *M. parishii* but with longer pistil. To determine which chromosome fragment was introgressed from *M. cardinalis* to *M. parishii* in the pistil NIL, we sampled 19 long-pistil individuals in the BC_2_S_1_ population (two rounds of backcrosses followed by one round of selfing) and performed bulked segregant analysis, following (Yuan *et al.*, 2016). Briefly, we pooled DNA samples from these 19 long-pistil individuals with equal representation from each sample. A small-insert (350-bp) library was prepared for the pooled sample, and 150-bp paired-end reads were generated by an Illumina NovaSeq platform at Novogene (Sacramento, CA), with a ~40-fold genome coverage. The resulting short reads were mapped to the *M. parishii* reference genome (http://mimubase.org/FTP/Genomes/Mparg_v2.0/) with CLC Genomics Workbench 7.0, and four chromosome fragments introgressed from *M. cardinalis* were identified (0-2.34 Mb on chromosome 1, 0-4.1 Mb on chromosome 4, 2.42 Mb -7.4 Mb on chromosome 5; and 0-51.76 Mb on chromosome 6). By genotyping the individual samples of the BC_2_S_1_ population using markers within the identified fragments, we found that the chromosome 4 and 6 fragments co-segregated with pistil length. That is, the pistil NIL resulted from phenotypic selection contains two causal loci introgressed from *M. cardinalis*. Genotyping a BC_3_S_1_ population further narrowed the chromosome 6 locus down to a genomic interval at 42.85 Mb-51.76 Mb.  Selfing a BC_3_S_1_ individual that is homozygous for *M. parishii* at the chromosome 1 and 5 fragments but heterozygous for the chromosome 4 and 6 fragments generated nine genotypes across the two loci, both co-segregating with pistil length. This also allowed us to decompose the BC_3_ NIL into two NILs.

A similar crossing approach was used to generate a CLL (flower size) NIL. To determine which chromosome fragment (s) was introgressed from *M. cardinalis* to *M. parishii* in the flower size NIL, we sequenced one BC_3_S_1_ individual most similar to *M. parishii* but with larger flower size. Based on deep sequencing, three heterozygous chromosome fragments (1.74 Mb-39 Mb on chromosome 3; 43.54 Mb-60.3 Mb on chromosome 6; 0-29.34 Mb on chromosome 7) and three *M*. *cardinalis* homozygous chromosome fragments (39 Mb-44.34 Mb on chromosome 3; 60.32 Mb-61.12 Mb on chromosome 6; 29.34 Mb-29.82 Mb on chromosome 7) were identified. Genotyping additional individuals of the BC_3_S_1_ population using markers located within the identified fragments revealed that only the chromosome 6/7 fragment (chromosomes 6 and 7 are linked together due to a reciprocal translocation) co-segregated with flower size.

**Sotola VA, Berg CS, Samuli M, Chen H, Mantel SJ, Beardsley PA, Yuan Y-W, Sweigart AL, Fishman L**. **2023**. Genomic mechanisms and consequences of diverse postzygotic barriers between monkeyflower species. *Genetics* **225**: iyad156.

**Yuan Y-W, Rebocho AB, Sagawa JM, Stanley LE, Bradshaw HD**. **2016**. Competition between anthocyanin and flavonol biosynthesis produces spatial pattern variation of floral pigments between Mimulus species. *Proceedings of the National Academy of Sciences of the United States of America* **113**: 2448–2453.
